# The effects of soil phosphorous content on microbiota are driven by the plant phosphate starvation response

**DOI:** 10.1101/608133

**Authors:** Omri M. Finkel, Isai Salas-González, Gabriel Castrillo, Stijn Spaepen, Theresa F. Law, Paulo José Pereira Lima Teixeira, Corbin D. Jones, Jeffery L. Dangl

**Author notes:** These authors contributed equally to this work (Omri M. Finkel, Isai Salas-González, Gabriel Castrillo).

## Abstract

Phosphate starvation response (PSR) in non-mycorrhizal plants comprises transcriptional reprogramming resulting in severe physiological changes to the roots and shoots and repression of plant immunity. Thus, plant-colonizing microorganisms – the plant microbiota – are exposed to direct influence by the soil’s phosphorous (P) content itself, as well as to the indirect effects of soil P on the microbial niches shaped by the plant. The individual contribution of these factors to plant microbiota assembly remains unknown. To disentangle these direct and indirect effects, we planted PSR-deficient Arabidopsis mutants in a long-term managed soil P gradient, and compared the composition of their shoot and root microbiota to wild type plants across different P concentrations. PSR-deficiency had a larger effect on the composition of both bacterial and fungal plant-associated microbiota composition than P concentrations in both roots and shoots. The fungal microbiota was more sensitive to P concentrations *per se* than bacteria, and less depended on the soil community composition.

Using a 185-member bacterial synthetic community (SynCom) across a wide P concentration gradient in an agar matrix, we demonstrated a shift in the effect of bacteria on the plant from a neutral or positive interaction to a negative one, as measured by rosette size. This phenotypic shift is accompanied by changes in microbiota composition: the genus *Burkholderia* is specifically enriched in plant tissue under P starvation. Through a community drop-out experiment, we demonstrate that in the absence of *Burkholderia* from the SynCom, plant shoots accumulate higher phosphate levels than shoots colonized with the full SynCom, only under P starvation, but not under P-replete conditions. Therefore, P-stressed plants allow colonization by latent opportunistic competitors found within their microbiome, thus exacerbating the plant’s P starvation.

## Introduction

Plant-derived carbon is the primary energy source for terrestrial heterotrophs, most of which are microbial. The interaction of these microbial heterotrophs with plants ranges between the extremes of mutualistic symbiosis [1] and pathogenesis [2,3]. However, the vast majority of plant-associated microbial diversity, the plant microbiota, lies between these two extremes, inducing more subtle, context-dependent effects on plant health [4–6]. The microbiota consumes plant photosynthate [7–9], and it provides benefits via protection from pathogens [10–14] or abiotic stress [15,16] or by increasing nutrient bioavailability [4,17,18].

The microbial community composition in soil, while governed by its own set of ecological processes [19], has an immense influence on the composition of the plant microbiota [20–22]. Correlations with soil microbial diversity, and by derivation, with plant microbiota composition and diversity, were observed for soil abiotic factors such as pH [19,23–25], drought [25–30] and nutrient concentrations [19,25,31–35]. Soil nutrient concentrations, in particular orthophosphate (Pi) – the only form of phosphorous (P) that is available to plants – produce comparatively modest to unmeasurable changes in microbial community composition [35,36]. Nevertheless, available soil Pi concentrations influences where a plant-microbe interaction lies along the mutualism-pathogenicity continuum [17].

Non-mycorrhizal plants respond to phosphate limitation by employing a range of phosphate starvation response (PSR) mechanisms. These manifest as severe physiological and morphological changes to the root and shoot, such as lateral root growth prioritization and depletion of shoot Pi stores [37]. In Arabidopsis, most of the transcriptional PSR driving these physiological responses is controlled by the two partially redundant transcription factors PHOSPHATE STARVATION RESPONSE 1 (PHR1) and PHR1-LIKE (PHL1) [38]. As a result, the double mutant *phr1 phl1* has an impaired PSR and accumulates a low level of Pi. Pi transport into roots relies on the PHOSPHATE TRANSPORTER TRAFFIC FACILITATOR1 (PHF1) gene, which is required for membrane localization of high-affinity phosphate transporters [39]. In axenic conditions, *phf1* mutants constitutively express PSR and accumulate low levels of Pi [39].

The plant’s response to its nutrient status is linked to its immune system. PHR1 negatively regulates components of the plant immune system, which can lead to enhanced pathogen susceptibility but also to the alteration of the plant’s microbiota under phosphate starvation [4]. Arabidopsis microbiota are altered in *phr1 phl1* and *phf1* mutants [4,36] in experiments using both natural and synthetic microbial communities [4].

Here, we examined (i) the effect of soil phosphorus (P) content on plant microbiota composition; (ii) how PSR modulates the plant microbiota and (iii) the interplay between PSR and soil P content in shaping the plant microbiota composition. We used a combination of greenhouse experiments with differentially P-fertilized soils, Arabidopsis PSR mutants and laboratory microcosms utilizing tractable synthetic bacterial communities. Using PSR mutants planted in P-amended soil, we demonstrate that the plant PSR regulators have a profound effect on the composition of root and shoot microbiota, overshadowing the effect of the soil P content. We constructed an ecologically tractable system utilizing a complex bacterial synthetic community (SynCom) as a model of the plant root microbiome and used this system to study the interactions between microbiota assembly and abiotic stress. We demonstrate deterministic responses of the SynCom members to changes in Pi concentrations, and we identify Pi-dependent plasticity along the mutualist-pathogen continuum.

## Results

### Phosphate starvation response in soil diverges from axenic *in vitro* assays

To better understand the effect of PSR genes on the plant microbiome under both Pi-limiting and Pi-replete conditions, we investigated how microbiota adapted to varying soil P levels interact with the plant’s PSR. We grew wild-type (wt) Arabidopsis and the PSR mutants *phf1* and *phr1 phl1* in soils collected from the ‘Halle long-term soil fertilization experiment’, ongoing since 1949 [40]. Each transect of soil has received one of three P fertilization treatments: zero (low), 15 (medium) and 45 (high) Kg[P].Ha^−1^. Year^−1^, resulting in a 3-5 fold difference in bioavailable P between the low and high treatments [41]. To differentiate the long-term adaptive effect of P limitation on the microbial community from the effect of short-term changes in P availability, we also fertilized a subset of the low P soil at the time of planting, and designated this condition low+P (Materials and Methods 1a-c, 4a).

We examined whether PSR, defined and typically studied in axenic conditions, is active in our soil-based experimental system. We harvested 8-week old plants grown in the different soils and quantified developmental and molecular phenotypes typically associated with PSR in both wt plants and mutants (Materials and Methods 1d, 1g). We found a strong positive correlation among all developmental features analyzed: shoot area, shoot fresh weight and shoot Pi accumulation across all soil conditions (Fig S1A and S1 Table). Shoot Pi accumulation showed the highest signal-to-noise ratio (Fig 1A, S1B and S1C Fig). In wt plants, shoot Pi accumulation was correlated with soil P conditions (Fig 1A). As expected [42], *phr1 phl1* showed a dramatic reduction in all phenotypic parameters (Fig 1A, S1B and S1C Fig) and *phf1* accumulated less shoot Pi than wt, but did not display any obvious morphological effect (Fig 1A and S1B-S1D Fig).

**Figure 1.**
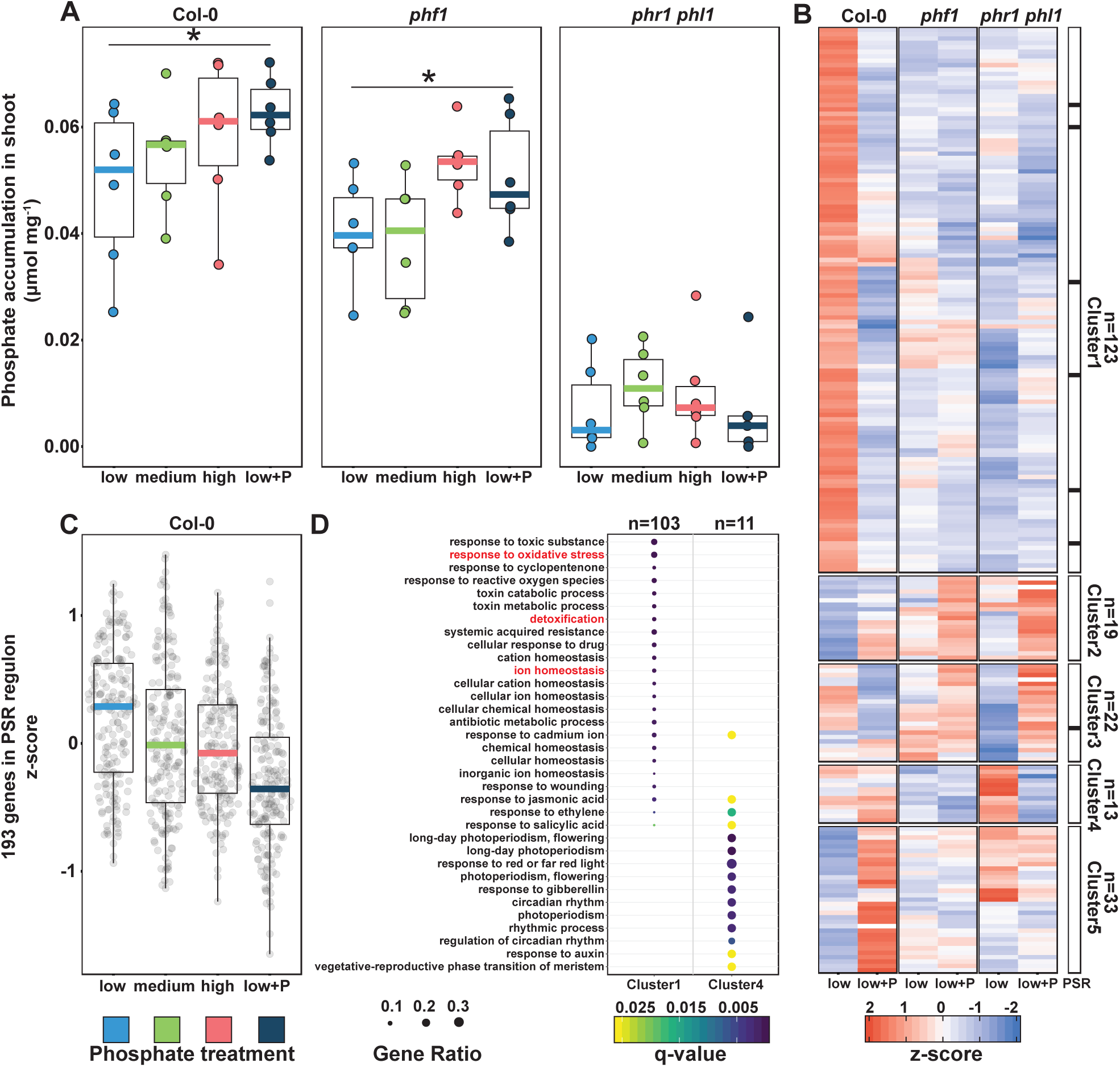
Plants respond to differential P conditions in soil. **(A)** Free phosphate content normalized by shoot fresh weight (mmol·mg-1) across wild type Col-0 plants and two PSR mutants, phf1 and phr1 phl1 (Materials and Methods 2a). Statistical significance between low P and low+P treatments was determined across each genotype independently by a paired t-test (p-value < 0.05). **(B)** Heatmap showing the average standardized expression of 210 differentially expressed genes (DEGs) across the low P and low+P samples in the Col-0, phf1 and phr1 phl1 genotypes (Materials and methods 4g). The black bar to the right highlights the distribution of seven genes belonging to the in vitro defined phosphate starvation response (PSR) marker genes [4] across the five clusters in the heat-map. **(C)** Average expression of 193 PSR marker genes [4] across the four phosphorus regimes in the Col-0 genotype (Materials and methods 4g). **(D)** Gene ontology (GO) enrichment for Clusters 1 and 4. Clusters 2, 3 and 6 did not show any statistically significant GO enrichment. The gene ratio is the proportion of genes per cluster that belong to a GO category.

To identify the transcriptomic signature of PSR in a low P soil, we compared the transcriptomes of the three genotypes from the low P samples to those of the low+P samples (Materials and Methods 1f, 3c, 4f-g and S2 Table). Using a likelihood ratio test (Materials and Methods 4g), we identified 210 genes that were differentially expressed across genotypes and P conditions (*q*-value < 0.1). After hierarchical clustering, 123 (59%) of these genes fall into a single cluster (cluster 1) of co-expressed genes that are exclusively highly expressed in wt under low P and whose expression requires *phr1 phl1* (Fig 1B). Thus, these genes represent a PSR under these experimental conditions. A gene ontology enrichment analysis (Fig 1D) illustrates that these genes are involved in processes such as ion homeostasis, detoxification and response to oxidative stress. Interestingly, few PSR genes defined from *in vitro* experiments were significantly differentially expressed in our soil experiment. From a previously defined set of 193 PSR marker genes defined using seven day old seedlings exposed *in vitro* to P limitation for up to two days [4], only seven were called as significant in our experiment using eight week old plants. Nevertheless, all seven of these genes were enriched in wt in low P soil (Fig 1B and S2 Table). Surprisingly, *phf1*, which was shown *in vitro* to constitutively express PSR genes, had a gene expression profile similar to *phr1 phl1* and did not exhibit the *phr1 phl1* dependent PSR response we observed for wt. This suggests a hitherto unknown link between PHF1 and the PSR-responsive genes seen under these experimental conditions. To corroborate that the canonical *in vitro*-defined PSR is also induced in wt plants, we compared the median expression of the set of 193 PSR marker genes [4] across the different soils (Fig 1C and S1E Fig). As expected, shoot Pi content was significantly correlated with the induction of PSR marker genes (Fig 1C and S1F Fig), with the highest median expression level in the low P conditions. We conclude that while the response of eight-week old plants to low P conditions in natural soil is markedly different from *in vitro*-defined PSR, wt plants indeed respond to low P conditions in the soils tested in a Pi concentration- and *phr1 phl1*-dependent manner.

### Bacterial and fungal plant microbiota differ in plant recruitment patterns

We studied the relationship between PSR and the plant microbiome in wild-type plants and the two PSR mutants grown in all four soils. Total DNA was extracted from shoots, roots and soil and the 16S (V3-V4) and ITS1 regions were amplified and sequenced to obtain bacterial and fungal community profiles, respectively. Bacterial sequences were collapsed into amplicon sequence variants (ASVs) while fugal sequences were clustered into operational taxonomic units (OTUs) (Materials and Methods 1e, 3a-b, 4b-c). Bacterial and fungal alpha- and beta-diversity measures conform to previously published data [20,36]: Microbial diversity decreased from the soil to the root and shoot compartments (Fig 2A, 2D, S2A, S2B Fig and S3 Table) and roots and shoots harbor bacterial and fungal communities distinct from the surrounding soil community and from each other (Fig 2B, 2E, S2C-S2F Fig and S4 Table). Plant-derived samples were primarily enriched in comparison to soil with members of the phyla Proteobacteria, Bacteroidetes and Actinobacteria and depleted in members of Acidobacteria and Gemmatimonadetes (Fig 2C, S2E Fig and S4 Table). Plant-enriched fungal OTUs belonged mainly to the phyla Ascomycota (orders Hypocreales and Pleosporales) and Basidiomycota (order Agaricales). Plant-depleted fungal OTUs belonged to Saccharomycetales (Ascomycota), Holtermanniales (Basidiomycota) and Mortierellales (Zygomycota) (Fig 2F, S2F Fig and S5 Table).

**Figure 2.**
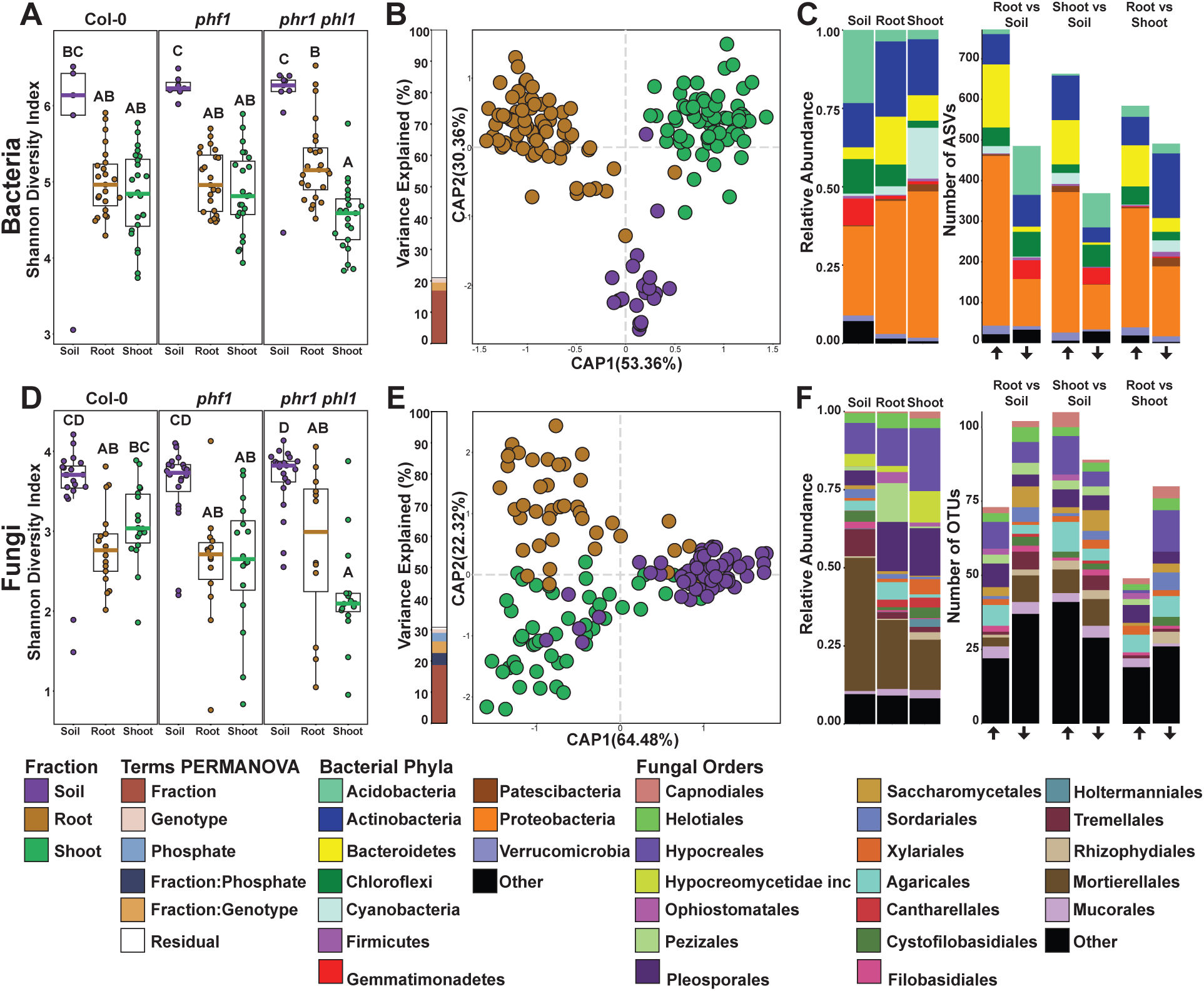
Plant recruitment patterns of bacteria and fungi. **(A,D)** Bacterial and fungal alpha diversity estimated using the Shannon Diversity Index (Materials and methods 4b). Letters represent post-hoc test results, based on a full factorial ANOVA model. **(B,E)** Canonical analysis of principal coordinates (CAP) based on Bray-Curtis dissimilarities between bacterial and fungal communities across the soil, root and shoot (Materials and methods 4b). The bar graph to the left of the CAP depicts the percentage of variance explained by statistically significant (p-value < 0.05) terms in a PERMANOVA model. **(C)** Left panel: Relative abundance profiles of the main bacterial phyla across the soil, root and shoot fractions. Right panel: Number of statistically significant amplicon sequence variants (ASVs) enriched in specific fractions (Materials and methods 4b). The arrows on the bottom of the panel denote the direction of the enrichment relative to the name of the contrast tested, the up arrow means enrichment in the left fraction of the contrast, whereas the down arrow means enrichment in the right fraction of the contrast (e.g. RootvsSoil, up arrow enriched in root relative to soil, bottom arrow enriched in soil relative to root). A detailed interactive visualization of the bacterial enrichment patterns across the multiple taxonomic levels can be found at (https://itol.embl.de/tree/1522316254174701551987253). **(F)** Left panel: Relative abundance profiles of the main fungal orders across soil, root and shoot fractions. Right Panel: Number of statistically significant operational taxonomic units (OTUs) enriched in specific fractions (Materials and methods 4b). The arrows on the bottom of the panel denote the direction of the enrichment relative to the name of the contrast tested, the up arrow signifies enrichment in the left fraction of the contrast, whereas the down arrow signifies enrichment in the right fraction of the contrast (e.g. RootvsSoil, up arrow enriched in root relative to soil, bottom arrow enriched in soil relative to root). A detailed interactive visualization of the fungal enrichment patterns across the multiple taxonomic levels can be found at (https://itol.embl.de/tree/1522316254174721551987262).

To quantify the effect of soil community composition on root and shoot microbiota composition, we used Mantel tests to detect correlation between dissimilarity matrices of the three fractions (root, shoot and soil). For bacteria, both root and shoot community dissimilarities were strongly correlated with soil (S3A and S3B Fig), while for fungi no correlation was detected with soil (S3D and S3E Fig). This observation indicates that the composition of both root and shoot communities are strongly dependent on soil community composition, despite the fact that bacterial microbiota are distinct from the soil community (Fig 2B). By contrast, the fungal microbiota composition both above and belowground is independent of the soil inoculum. This difference implies that the plant’s microbiota filtering mechanisms are fundamentally different for fungi and bacteria.

### Shoot and root microbiota are both correlated and distinct

Shoot and root samples are rarely analyzed in the same study [43]. We show here that roots and shoots harbor distinct communities from each other (Fig 2B, 2E, S2C-S2F Fig and S4 Table). To further explore organ specificity in the plant microbiome composition, we compared root and shoot samples at the OTU/ASV level. Shoots were mainly enriched with the bacterial phyla Cyanobacteria and Patescibacteria compared to the root, while roots were enriched with Proteobacteria, Chloroflexi and Bacteroidetes (Fig 2C, S2E Fig and S4 Table). With regard to fungal orders, shoots were enriched with Capnodiales, Hypocreales and Rhizophydiales while roots were enriched with Pezizales, Xylariales and Mucorales (Fig 2F, S2F Fig and S5 Table). The shoot enrichment of Cyanobacteria points to the availability of light as an important factor in niche differentiation within the plant [44–46]. We used Mantel tests to detect correlation between dissimilarity matrices of root and shoot samples. Despite the fact that they harbor distinct communities, roots and shoots were correlated with each other for both bacteria and fungi (S3C Fig and S3F Fig). Thus, while roots and shoots form distinct bacterial and fungal niches, shifts in microbiota in both of these niches are correlated, suggesting co-inoculation between plant fractions.

### The plant microbiome composition is driven by the plant PSR status

We investigated the influences of plant PSR signaling and the different soil P concentrations on microbial community composition (Materials and Methods 4c). Constrained ordination showed significant differences between both bacterial and fungal communities across the P accumulation gradients represented by the different soils and PSR mutants (Fig 3A, 3B and S4A, S4B Fig). For both bacteria and fungi, the effect of PSR status in roots was stronger than the soil P effect (Fig 3A and 3B). Notably, the effect of PSR status on the bacterial community overrides the effect of soil P (Fig 3A and S4A Fig), whereas for the fungal community, both plant PSR and soil P had a strongly significant effect (Fig 3B and S4B Fig). This is contrary to previous results from the same soils where there was no detectable P effect on microbial community composition [41]. In shoots, both bacteria and fungi responded to PSR genotype, but in this case, did not respond to soil P (S4C Fig and S4D). We did not observe a significant soil P:genotype interaction effect for either bacteria or fungi (S4A and S4B Fig), confirming that *phf1* and *phr1 phl1* both had atypical bacterial microbiomes regardless of Pi status. As expected, we did not observe a PSR effect in the soil samples (S4E and S4F Fig).

**Figure 3.**
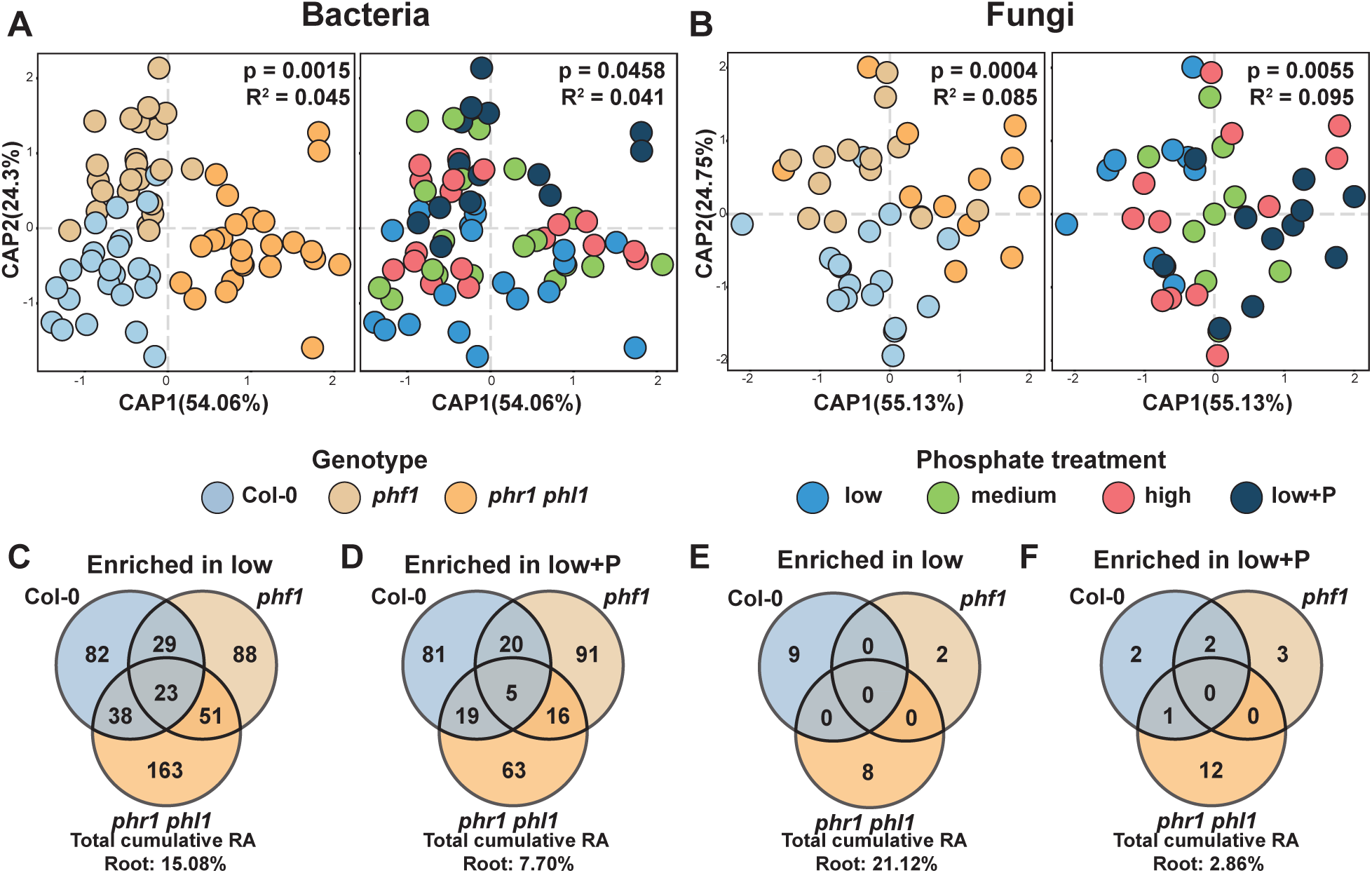
Plant phosphate starvation response controls the assembly of the plant microbiome. **(A,B)** Canonical analysis of principal coordinates showing the influence of plant genotypes and soil phosphorus content over the **(A)** bacterial and **(B)** fungal communities in the root (Materials and methods 4b). The p-value and R2 values inside each plot are derived from a PERMANOVA model and correspond to the genotype and phosphorus term respectively. **(C,E)** Venn diagrams showing the distribution of **(C)** bacterial ASVs and **(E)** fungal OTUs with statistically significant (q-value < 0.1) higher abundance in the low P treatment in comparison to the low+P treatment in the Col-0, phf1 and phr1 phl1 roots (Materials and methods 4b). **(D,F)** Venn diagrams showing the distribution of **(D)** bacterial ASVs and **(F)** fungal OTUs with statistically significant (q-value < 0.1) higher abundance in the low+P treatment in comparison to the low P treatment across the Col-0, phf1 and phr1 phl1 roots. RA=relative abundance.

The relatively weak soil P effect observed here indicates that most of the soil effect on bacterial microbiota mentioned above (S3A and S3B Fig) should be explained by other edaphic factors. On the other hand, the notable genotype effect illustrates that the plant niche filtering (Fig 2B and 2E) is partly shaped by PSR.

To define which taxa at the ASV (Bacteria) and OTU (Fungi) levels were influenced by soil P and/or plant PSR, we applied a generalized linear model (GLM, Materials and Methods 4c) to the count datasets, contrasting the low P samples against the low+P samples. We detected 769 bacterial ASVs (S6 Table) and 39 fungal OTUs (S7 Table), accounting for 23% and 24% of the bacterial and fungal abundance in the root, respectively, that were differentially abundant in at least one genotype (Fig 3C-3F). Of these, 568 bacterial ASVs and 36 of the fungal OTUs were genotype-specific, suggesting these taxa respond to PSR-regulated processes, rather than the P concentration in the soil.

Taken together, these results indicate that plant microbiota are relatively robust to differences in soil P content, but are sensitive to the plant PSR status. Responses to soil P concentration are contingent on PSR regulatory elements under both low and high P conditions.

### Bacterial synthetic communities modulate the plant PSR

The results obtained from the soil experiment suggest that niche sorting in the plant microbiome is not only determined by first order interactions (plant-microbe, microbe-microbe, microbe-environment), but also by higher-order interactions, such as the effect of abiotic conditions on plant-microbe interactions. This is evident in the large proportion of ASVs/OTUs that respond to soil P in a genotype-specific manner (Fig 3C-3F). To establish a system where interactions of different orders of complexity can be studied reproducibly, we constructed a plant-microbe microcosm that can be deconstructed to its individual components, while retaining a complexity that is comparable to natural ecological communities. We designed a representative bacterial synthetic community from a culture collection composed of isolates derived from surface-sterilized Arabidopsis roots [47] (Materials and Methods 2a). We selected 185 genome-sequenced isolates representing a typical plant-associated taxonomic distribution (Fig 4A and 4B). We grew each isolate separately and mixed the grown cultures to equal optical densities. We grew 7-day-old Arabidopsis seedlings in a Pi concentration gradient (0, 10, 30, 50, 100, 1000 µM KH_2_PO_4_) and concomitantly exposed them to the bacterial SynCom on vertical agar plates for 12 days (Materials and Methods 2c-d).

**Figure 4.**
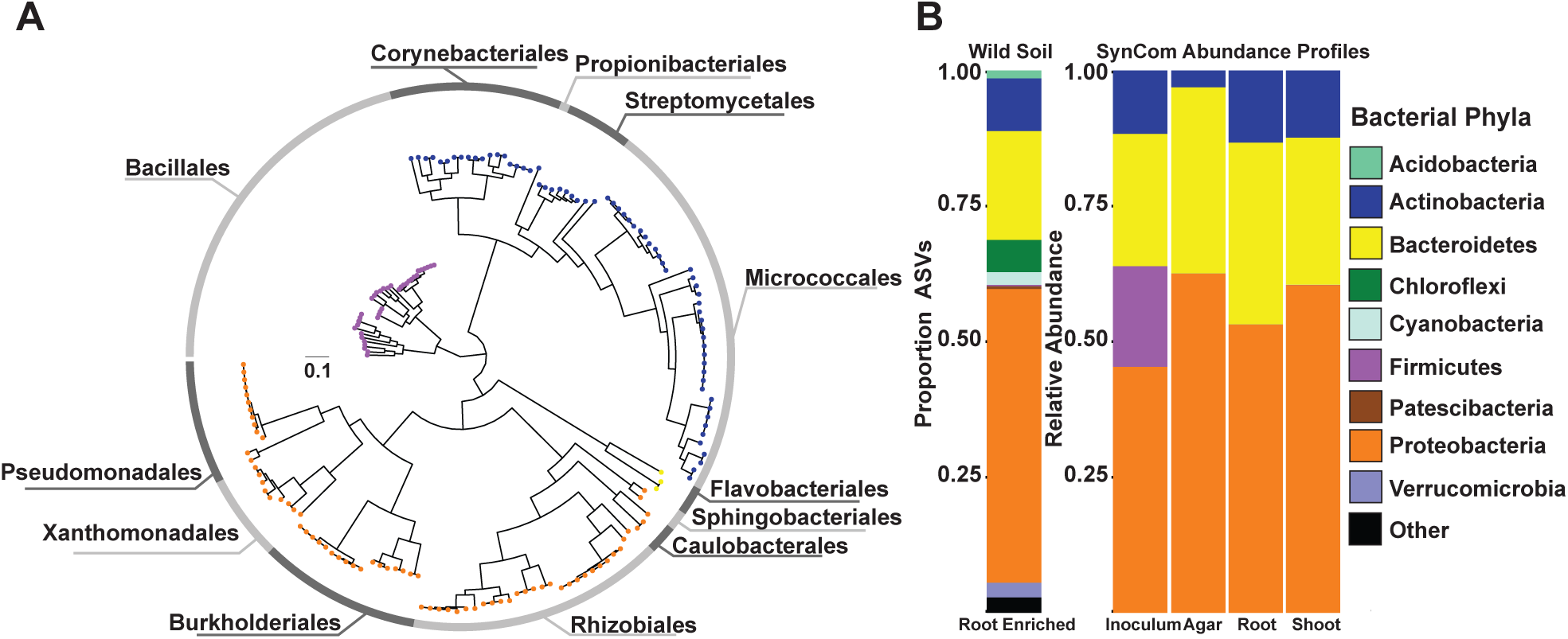
Bacterial synthetic community reproduces the typical plant-associated taxonomic distribution found in soil. **(A)** Phylogenetic tree of 185 bacterial genomes included in the synthetic community (SynCom) (Materials and methods 4e). The tree tips are colored according to the phylum classification of the genome in **(B)**, the outer ring shows the distribution of the 12 distinct bacterial orders present in the SynCom. **(B)** Left Panel: Proportion of amplicon sequence variants (ASVs) enriched in the root in comparison to the natural soil across all treatments and genotypes based on a fitted generalized linear model (q-value < 0.1). Each ASV is colored according to its phylum level classification. Right Panel: Relative abundance profiles of bacterial isolates across the initial bacterial inoculum, planted agar, root and shoot fractions. Each isolate is colored according to its phylum level classification based on the genome-derived taxonomy.

First, we investigated whether PSR is induced in our experimental system. Similar to the natural soil-based experiment, we quantified developmental and transcriptional phenotypes associated with PSR in plants grown in different concentrations of Pi (Materials and Methods 2f-g). The presence of the SynCom consistently decreased root length across all Pi concentrations, but the Pi gradient did not affect root length (S5A Fig). Shoot size was correlated with Pi concentration, and the slope of this trend was affected by the presence of the SynCom: at High Pi the SynCom tended to increase shoot size, while at low Pi the SynCom decreased it (Fig 5A), suggesting that the microbiome plays a role in shaping the plant’s response to different Pi concentrations.

**Figure 5:**
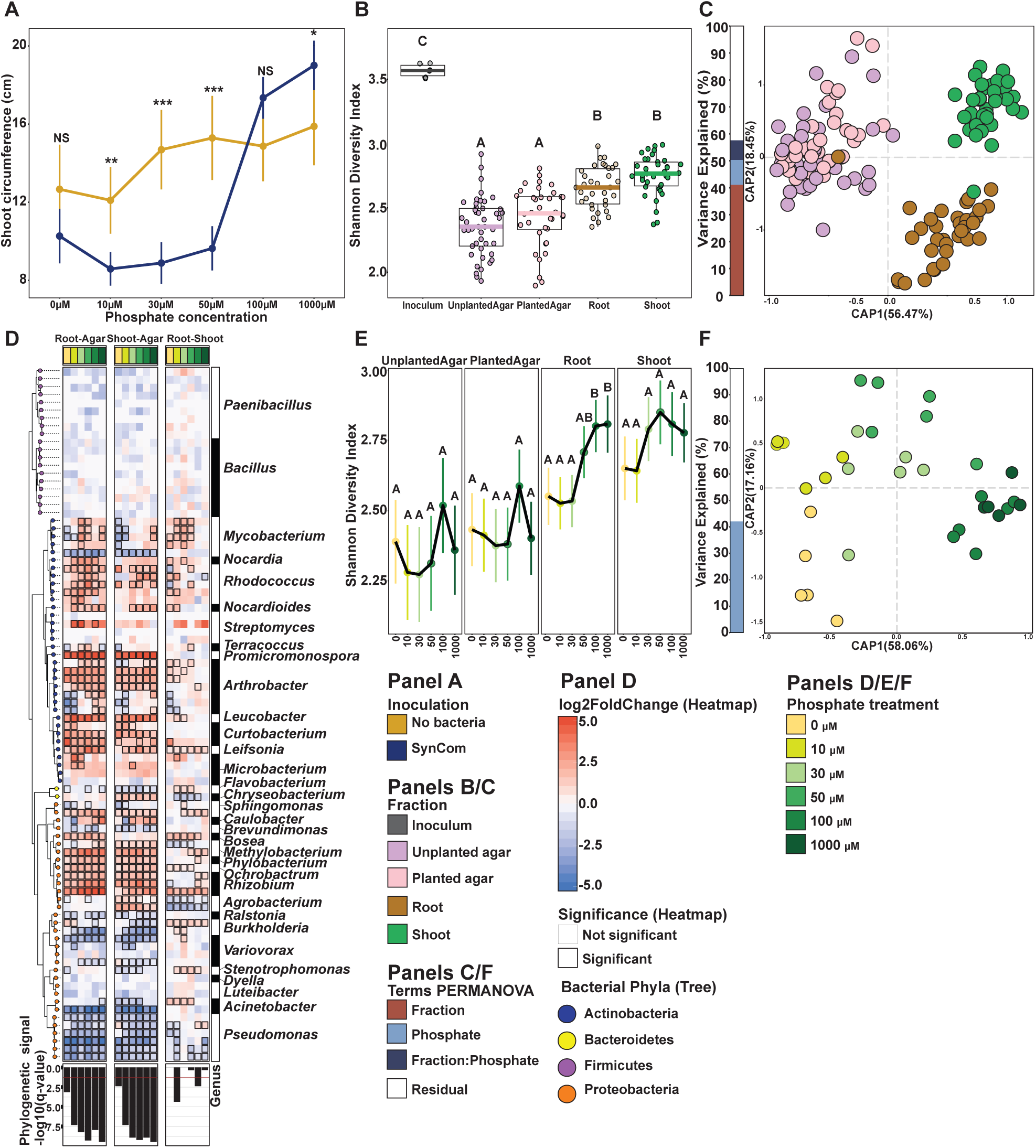
Synthetic bacterial communities display deterministic niche sorting in plants. **(A)** Stripchart displaying the average shoot size of plants grown across a Pi gradient either in sterile conditions or with the SynCom (Materials and methods 2g). Each dot in the scatterplot represents the mean value for that particular treatment, the range crossing each dot represents the 95% confidence interval calculated. The lines are drawn to connect the means. **(B)** Alpha diversity across the fractions sampled was estimated using the Shannon Diversity index (Materials and methods 4d). An ANOVA model followed up by a Tukey HSD test were applied to estimate differences between inoculum, unplanted agar, planted agar, root and shoot fractions. Lefters represent the results of the post hoc test. **(C)** Canonical analysis of principal coordinates (CAP) based on Bray Curtis dissimilarities between bacterial communities across the four fractions sampled (Materials and methods 4d). The bar graph to the left of the CAP depicts the percentage of variability explained by statistically significant (p-value < 0.05) terms in the PERMANOVA model. **(D)** Enrichment patterns of the SynCom (Materials and methods 4d). Each row along the different panels of the figure represents a USeq: a USeq encompasses a set of indistinguishable V3-V4 16S rRNA sequences present in the 185-member-SynCom. Phylogenetic tree (on the left) is colored based on the phylum-level classification of the corresponding USeq. Each column in the heatmaps represents a specific contrast in the enrichment model. We calculated root vs agar (left heatmap), shoot vs agar (middle heatmap) and root vs shoot (right heatmap) enrichments within each Pi treatment (e.g. Root_0Pi vs Agar_0Pi, Materials and methods 4d). The heatmaps are colored based on log2 fold changes derived from the fitted GLM. Positive fold changes (colored in red gradient) represent enrichments on the left side of the name of the contrast (e.g. Root-Agar, enriched in root in comparison to agar), whereas negative fold changes (colored in blue gradient) represent enrichments on the right side of the name of the contrast (e.g. Root-Agar, enriched in agar in comparison to agar). The bottom panel depicts the transformed (-log10) q-value derived from a phylogenetic signal Pagel’s λ test. Tests were performed per column in the heatmap (e.g. Root0µM pi vs Agar0µM pi). **(E)** Bacterial alpha diversity estimated using the Shannon Diversity index (Materials and methods 4d). Lefters represent the results of the post hoc test. Lines connect the means. **(F)** Canonical analysis of principal coordinates showing the influence of phosphate on the bacterial communities in the root (Materials and methods 4d). The bar graphs to the left of the CAP depict the percentage of variability explained by statistically significant (p-value < 0.05) variables based on a PERMANOVA model.

We performed RNA-Seq on inoculated and uninoculated plants exposed to high (1000 µM) and low (50 µM) Pi (Materials and Methods 3c, 4h). To confirm that our low Pi treatments induce PSR, we examined the expression of the 193 PSR markers defined in [4]. We found that 168 of the 193 PSR markers genes were significantly induced in uninoculated plants at low Pi as compared with high Pi conditions. In the presence of the SynCom, 184 out of 193 PSR marker genes were significantly induced, and the average fold change increased from 4.7 in uninoculated conditions to 11 in the presence of the SynCom (S5B Fig). We further examined whether the 123 low P responsive genes from the soil experiment (Cluster 1 in Figure 1B) are overexpressed in the agar system as well. We found that 59 of the 123 genes (47.2%) were low Pi-enriched in uninoculated plants and 72 (58.5%) were low Pi-enriched in the presence of the SynCom. The average fold change for this set of 123 genes was 1.6 in uninoculated conditions and 2.0 in the presence of the SynCom (S5C Fig). These results confirm that (i) in both our systems, wild soils and axenic conditions, PSR is induced at low Pi and (ii) the SynCom enhances this induction, similar to the results reported in [4].

### Bacterial synthetic communities display deterministic niche sorting in the plant microbiome

To quantify the establishment of the SynCom in the plants, we determined bacterial community composition after 12 days of co-inoculation in roots, shoots and agar via 16S rRNA gene amplicon sequencing, mapping reads to 97 unique sequences (USeq) representing the 185-strain SynCom (Materials and methods 2e, 3a, 4d-e, S8 Table). We found that plant roots and shoots sustained a higher bacterial alpha diversity than the surrounding agar (Fig 5B, S9 Table), an aspect in which our experimental system differs from a natural environment where species richness is higher in the surrounding soil than in the plant (Fig 2A). As in natural soil experimental systems, agar, roots and shoots assembled distinct bacterial communities and this difference among these three fractions explained most of the variance in community composition despite the different Pi-concentrations (Fig 5C and S5D Fig).

To study which strains are enriched in the root and shoot under the different Pi concentrations, we utilized a GLM (S10 Table, Materials and Methods 4d). Noticeably, plant (root and shoot) enrichment is strongly linked to phylogeny (Fig 5D) and is robust across the phosphate gradient assayed. In contrast, the root vs shoot comparison did not exhibit a significant phylogenetic signal, highlighting the fact that the ability to differentially colonize the shoot from the root under these conditions is phylogenetically scattered across the SynCom. As in the soil census, shoot, root and agar beta diversities were significantly correlated (S5E-S5G Fig).

We hypothesized that by establishing a standardized protocol for producing the inoculum and controlling the growth conditions, we will have created a reproducible, controlled system that prioritizes niche sorting over stochastic processes. To test this, we compared the amount of variance explained by a GLM in the natural community experiment vs the SynCom experiment. Supporting our hypothesis, only 1,518 out of 3,874 measurable ASVs (32% of the total ASVs), accounting for 72% of the relative abundance in plant tissue, shift significantly between root and soil in the natural community survey, while 58 out of 97 USeqs (59%), accounting for 99% relative abundance in plant tissue, were significantly enriched or depleted in plant tissue in the SynCom experiment (Fig 5D). This difference in tractability between the soil and microcosm experiment is also evident in the PERMANOVA model results: in the soil experiment, we could explain 21% of variance in bacterial community composition (Fig 2B, and S2C Fig), whereas in the microcosm, we could explain 57% of variance (Fig 5C and S5D Fig).

These results indicate that plant colonization is largely deterministic in our SynCom system, as opposed to microbiomes in nature, which are strongly driven by stochastic and neutral processes of community assembly [48]. The reproducibility of this system, coupled with our ability to edit it as a tool for hypothesis-testing, is crucial to bridge ecological observation with mechanistic understanding of plant-microbiota interactions.

### Phosphate stress-induced changes in the root microbiome

The shifting role of the SynCom from increasing shoot size under replete Pi to decreasing shoot size and PSR induction under Pi limitation (Fig 5A and S5B Fig) can be explained by either a shift in the lifestyle of individual bacteria along the mutualist-pathogen continuum or by changes in the microbiota composition along the Pi gradient. The latter would favor the proliferation of mutualist bacteria only when sufficient nutritional requirements are met. To measure the effect of the Pi concentration in the media on the SynCom composition in wt plants, we measured alpha and beta diversity along our Pi gradient (0, 10, 30, 50, 100, 1000 µM KH_2_PO_4_) in roots, shoots and agar. We observed a positive correlation between alpha diversity and Pi concentrations, resembling a partial ecological diversity-productivity relationship – the prediction/observation of a bell-shaped response of ecological diversity to environmental productivity [49,50] – in roots and shoots, but not in the surrounding agar (Fig 5E). As for beta diversity, the composition of the SynCom shifted significantly along the Pi concentration gradient (Fig 5F and S6A-S6E Fig). Pi-stressed plants therefore assemble an altered microbiome, shifting from a net-positive outcome for the plant, to a net-negative one, as measured by shoot size (Fig 5A).

### *Burkholderia* respond to Pi stress-induced changes in the plant

In a previous publication [4], we demonstrated that PHR1 negatively regulates defense-related genes under low-Pi conditions. Suppression of plant defense and consequent alterations in colonization could account for some of the shift we observed from a beneficial to a detrimental community. We thus aimed to identify bacteria that respond to Pi stress-induced changes in the plant, rather than the Pi concentration itself. To do so, we searched for USeqs that displayed a strong Pi:fraction (shoot, root, agar) interaction in our GLM (S7A Fig, S11 Table and Materials and Methods 4d). Two of the three USeqs displaying the strongest Pi:fraction interaction belonged to *Burkholderiaceae*, representing all 5 *Burkholderia* strains used in this experiment. The relative abundance of these USeqs is positively correlated with Pi concentration in the agar, but is negatively correlated with Pi concentration in the root and shoot (Fig 6A and S7B Fig). This pattern suggests that these strains are responding to physiological changes in the plant – either via a suppression of an immune mechanism that keeps them in check under high Pi, or via an unknown positive selection mechanism under low Pi.

**Figure 6.**
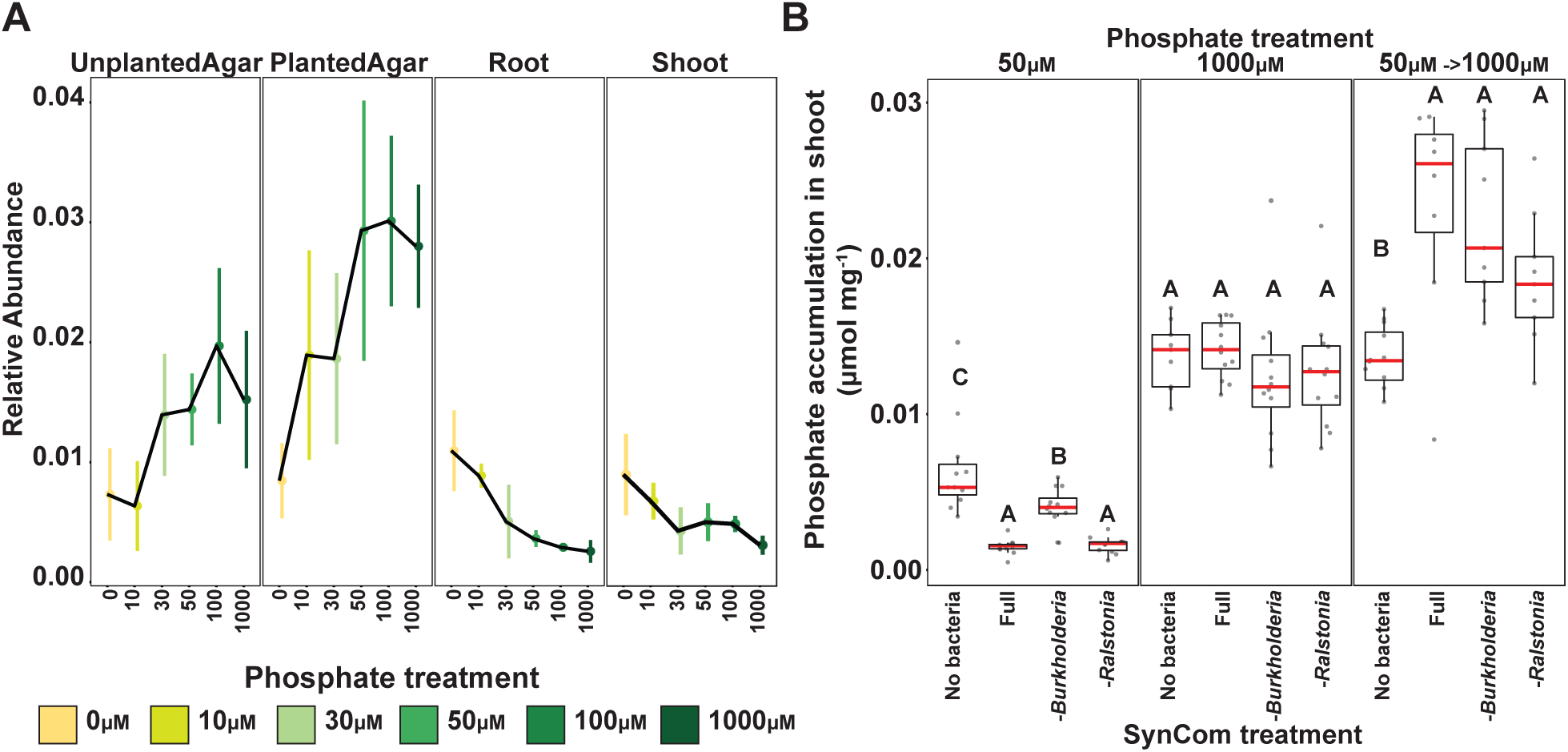
Bacterial strains respond to Pi-stress-induced physiological changes in the wild type plants. **(A)** Relative abundance of Burkholderia Useq 16 that exhibits a statistically significant (q-value < 0.1) Pi-enrichment between the plant fractions and the agar fraction (Materials and methods 4d). The middle dot of each strip bar corresponds to the mean of that particular condition, the range of the strip bar corresponds to the 95% confidence interval of the mean. The lines are drawn connecting the means for each Pi concentration. **(B)** Boxplots showing the phosphate accumulation in plants exposed to different synthetic communities across three phosphate treatments. Statistically significant differences among SynCom treatments were computed inside each phosphate treatment separately using an ANOVA model. Lefters represent the results of the post hoc test.

To measure the physiological effect of the specific recruitment of *Burkholderia* under Pi stress on the plant, we conducted a drop-out experiment in which we compared plants inoculated with the full SynCom to plants inoculated with a SynCom excluding all five *Burkholderia* isolates. We also included a SynCom excluding all members of the neighboring *Ralstonia* clade (Fig 4A), which didn’t display any discernible Pi-response. We measured Pi concentrations in the shoots (a proxy for PSR) of plants grown in high (1000 µM) and low (50 µM) Pi with the different SynComs. In addition, we measured shoot Pi in a re-feeding treatment with SynCom-inoculated plants grown in low (50 µM) Pi and then transferred to high Pi (1000 µM) conditions. All SynCom treatments decreased shoot Pi content in the low Pi conditions compared to the uninoculated plants but recovered to a higher shoot Pi level than the uninoculated treatments upon transferring to high Pi conditions, reproducing our previous report [4] (Fig 6B). Among inoculated treatments, plants colonized with the *Burkholderia* drop-out treatment (SynCom excluding all *Burkholderia*) had a higher Pi content than either plants colonized with the full SynCom or with the *Ralstonia* drop-out SynCom only in the low Pi conditions. There was no difference in shoot Pi among the SynCom treatments in either the high Pi treatment, or following the refeeding treatment. This finding suggests that the enrichment of *Burkholderia* in plant tissue under Pi starvation can be considered a shift in the effect of bacteria on the plant from a positive interaction to a negative one. This supposition is consistent with the plant immune system gating this taxon under replete Pi but being unable to do so under PSR.

## Discussion

Despite the fact that phosphate is a critical nutrient for plants and their microbiota, differences in phosphate content have relatively subtle effects on plant and soil microbiome compositions compared to abiotic factors like pH or drought, which cause pronounced, phylogenetically consistent changes in community configurations [19,27,29]. Several studies link host physiological response to the soil phosphate status with the bacterial [4,51] and fungal [34,36] microbiome. A recent report of Arabidopsis planted in a 60-year-long annual phosphorus fertilization gradient (the same soil used in the current study) showed a modest P effect on plant microbiome composition [41]. Previously, we showed that PSR mutants in Arabidopsis have subtly different bacterial microbiomes in Pi replete [4] conditions and a recent publication showed that PSR mutants had a slightly altered fungal microbiome in Pi replete but not in Pi depleted conditions [36].

Here, we analyzed fungi and bacteria side by side and demonstrated a pronounced effect of PSR impairment on both bacterial and fungal components of the plant microbiota. We noted an intriguing difference that emerged in the patterns of niche sorting between bacteria and fungi (S3A S3B, S3D and S3E Fig). The bacterial microbiota composition is strongly dependent on the soil bacterial community composition, whereas changes to the fungal microbiota are uncoupled from changes to the soil fungal community composition. This indicates that the plant is markedly more selective as to the fungi allowed to proliferate in its tissue than it is with bacteria. Similar to [36], amendment of the soil with P at the time of the experiment (low P vs low+P) caused a shift in the microbial community, albeit weaker than the effect of knocking-out PSR genes. Our results show that impairment of PSR genes profoundly affects the composition of the plant microbiota, under a range of P conditions, and that observed shifts in root-derived microbial communities may not be a result of sensitivity to P concentrations, but rather a response to PSR regulation in the hosts. This raises the alternative hypotheses that PSR-regulated shifts in microbiota composition are either adaptive to the plant, or reflect opportunistic strategies by bacteria, exploiting the repression of immunity by PSR regulation [4,17]. Under the former hypothesis microbes recruited by the plant under Pi stress provide the plants with an advantage vis a vis coping with this stress, whereas under the latter, opportunistic microbes might be making a bad situation worse for the plant. In the case of *Burkholderia* in our SynCom, results support the latter hypothesis. *Burkholderia* contribute to depletion of shoot Pi stores, only under Pi-limiting conditions. However, plant-adaptive microbial recruitment under low Pi has been shown to occur as well [17]. The fact that in soil bacteria responding to PSR genes are not a monophyletic group indicates that multiple mechanisms are involved. It is likely that these mechanisms encompass both plant-adaptive and opportunistic strategies.

The genus *Burkholderia* emerges as a PSR-responsive taxon. We examined the effect of *Burkholderia* on shoot Pi accumulation, which we’ve shown to be a reliable marker for PSR (S1F Fig). We compared the effect of *Burkholderia* on shoot Pi accumulation from within a full SynCom (a realistic proxy for the bacterial community) to that of the full SynCom lacking *Burkholderia*, a strategy akin to knocking-out a gene of interest, also recently applied in [52]. The control treatment for this type of approach is the full SynCom, while in a plant-bacterium binary association experiment it would typically be sterile conditions. As both sterile conditions and binary association are strong deviations from conditions that may be encountered in the field, the results of binary association experiments may be correspondingly distorted. Using the drop-out approach, we expect to see more subtle differences, as the microbial load on the plant doesn’t change much, but also that these differences be more relevant to the field; an expectation that is yet to be empirically tested. Our observation that dropping *Burkholederia* out of the SynCom increased shoot Pi in Pi limiting conditions (50 µM Pi) but not in Pi replete conditions (1000 µM Pi) suggests that strains in this genus shift their relationship with the plant from a seeming commensal to a competitor/pathogen. It is likely that this shift is related to the repression of plant immune function by key regulators of PSR in low Pi [4], suggesting a specific plant-dependent trade off during PSR.

This study shows that despite 60 years of differential fertilization, differences in PSR and in microbiome composition between the low P and high P soils are subtle, possibly because Pi status is highly buffered by the plant ionomic regulatory network [53]. Only when comparing the low P vs the P-supplemented low+P samples, is there a discernible difference in PSR (Fig 1A), which correlates to a stronger effect on microbiota composition. This suggests that bioavailable Pi added to the soil is quickly consumed, and short-term amendments are needed in order to detect changes. It is easier to produce Pi-limiting conditions *in vitro* using defined media, as evidenced from shifts in both PSR gene expression and microbiome composition in the microcosm system introduced here. Our SynCom, comprising 185 genome-sequenced endophytic bacterial isolates, was designed to resemble a natural bacterial community (Fig 4B). The community assembly patterns shown for this system are highly reproducible, demonstrating that microbiome assembly is largely a deterministic process. The reproducibility and editability of this system can be used for detailed mechanistic study of the processes that determine community assembly and its influence on plant phenotype and fitness.

## Materials and Methods

### 1. Soil P gradient experiment

#### a. Collection of soil from field site

Soil used in this experiment was collected from the long-term Pi fertilization field (“Field D”) trial at the Julius Kühn Experimental Station at Martin Luther University of Halle-Wittenberg (51°29′45.6′′N, 11°59′33.3′′E) [40,54]. Soil cores (10 cm diameter × 15 cm depth) were taken from 18 6 × 5 m unplanted plots, belonging to two strips. These plots represent three P fertilization regimens: low, medium and high P (0, 15 and 45 kg P ha^−1^ year^−1^, respectively). Strips were harvested independently in the middle of March (strip 1) and beginning of April (strip 2). Approximately 2 cm of the topsoil was discarded and the remaining lower 13 cm of soil was stored at 4 °C until use. Soils were homogenized with a mesh sieve wire (5 × 5 m^2^) and about 300 g of soil were added to each pot (7 × 7 × 7 cm^3^).

#### b. Experimental design

Each of the three Arabidopsis genotypes was grown in soil from all 18 plots (6 plots per P treatment). In addition, a 4^th^ P regimen designated ‘Low+P’ was created by adding additional P to a set of pots with low P. The amount of P added to these pots is based on the difference in total P between Low and High P plots. The average difference between Low and High P over all the plots is 42 mg P per kg soil [41]. Per pot, this is 12.6 mg P (accounting for 300 g soil per pot). Thus, a 10 ml solution consisting of 4.2 mg P in the form of 20% K_2_HPO_4_ and 80% KH_2_PO_4_ was added to the pots in 3 applications (Week 2, 4 and 6) before watering (in order to distribute the P through the soil).

Thus, the experiment included four soil treatments (low P, medium P, high P, low+P) and three genotypes (Col-0, *phf1* and *phr1 phl1*) with 6 independent replicates, amounting to 72 pots. Pot positions in the greenhouse was randomized.

#### c. Plant growth conditions

*Arabidopsis thaliana* ecotype Col-0 and mutants *phf1 and phr1 phl1* (both in the Col-0 background) were used. Seeds were surface sterilized (20 min 70% EtOH, 10 s 100% EtOH) and planted directly onto moist soil. Sown seeds were stratified for 3 days at 4 °C before being placed in a greenhouse under short-day conditions (6/18 day-night cycle; 19 to 21 °C) for 8 weeks. Germinating seedlings were thinned to four plants per pot.

#### d. Sample harvest

After eight weeks of growth, pots were photographed, and shoot size was quantified using WinRhizo software (Regent instruments Inc. Québec, Canada). For DNA extraction, two roots, two shoots and soil from each pot were harvested separately. Roots and shoots were rinsed in sterile water to remove soil particles, placed in 2 ml Eppendorf tubes with 3 sterile glass beads, then washed three times with sterile distilled water to remove soil particles and weakly associated microbes. Root and shoot tissue were then pulverized using a tissue homogenizer (TissueLyser II; Qiagen) and stored at −80 °C until processing. Five ml of soil from each pot was suspended in 20 ml of sterile distilled water. The resulting slurry was sieved through a 100 µm sterile cell strainer (Fisher Scientific) and the flow-through was centrifuged twice at maximum speed for 20 minutes, removing the supernatant both times. The resulting pellet was stored at −80 °C until processing. For RNA extraction, one root system and one shoot were taken from three replicates of each treatment, washed lightly to remove soil particles, placed in 2 ml Eppendorf tubes with three glass beads and flash frozen with liquid nitrogen. Tubes were stored at −80 °C until processing. For shoot Pi measurement, 2-3 leaves from the remaining shoot in each pot were placed in an Eppendorf tube and weighed. 1% acetic acid was then added and samples were flash frozen and stored at −80 °C until processing.

#### e. DNA extraction

DNA extractions were carried out on ground root and shoot tissue and soil pellets, using the 96-well-format MoBio PowerSoil Kit (MoBio Laboratories; Qiagen) following the manufacturer’s instruction. Sample position in the DNA extraction plates was randomized, and this randomized distribution was maintained throughout library preparation and sequencing.

#### f. RNA extraction

RNA was purified from plant tissue using the RNeasy Plant Mini Kit (Qiagen) according to the manufacturer’s instructions and stored at −80 °C.

#### g. Quantification of plant phenotypes

The Ames method [55] was used to determine the phosphate concentration in the shoots of plants grown on different Pi regimens and treatments. Shoot area was measured using WinRhizo software (Regens Instruments Inc.).

### 2. Bacterial SynCom experiment

#### a. Bacterial isolation and culture

The 185-member bacterial synthetic community (SynCom) contained genome-sequenced isolates obtained from *Brassicaceae* roots, nearly all Arabidopsis, planted in two North Carolina, USA, soils. Since both bacteria and fungi responded similarly to PSR in our soil experiments, we only included bacteria, which are more compatible with our experimental system, in our SynCom. A detailed description of this collection and isolation procedures can be found in [47]. One week prior to each experiment, bacteria were inoculated from glycerol stocks into 600 µL KB medium in a 96 deep well plate. Bacterial cultures were grown at 28°C, shaking at 250 rpm. After five days of growth, cultures were inoculated into fresh media and returned to the incubator for an additional 48 hours, resulting in two copies of each culture, 7 days old and 48 hours old. We adopted this procedure to account for variable growth rates of different SynCom members and to ensure that non-stationary cells from each strain were included in the inoculum. After growth, 48-hour old and 7-day old plates were combined and optical density (OD) of the culture was measured at 600 nm using an Infinite M200 Pro plate reader (TECAN, Switzerland). All cultures were then pooled while normalizing the volume of each culture according to the OD (we took a proportionally higher volume of culture from cultures with low OD). The mixed culture was then washed twice with 10 mM MgCl_2_ to remove spent media and cell debris and vortexed vigorously with sterile glass beads to break up aggregates. OD of the mixed, washed culture was then measured and normalized to OD=0.2. 100 µL of this SynCom inoculum was spread on each agar plate prior to transferring seedlings.

#### b. Experimental design of agar experiments

We performed the Pi-gradient experiment in two independent replicas (experiments performed at different time, with fresh bacterial inoculum and batch of plants), each containing three internal replications, amounting to six samples for each treatment. We had two SynCom treatments: no bacteria (NB) and SynCom; six Pi concentrations: 0, 10, 30, 50, 100 or 1000 µM Pi; and two plant treatments: planted plates, and unplanted plates (NP).

For the drop-out experiment, the entire SynCom, *excluding* all five *Burkholderia* and both *Ralstonia* isolates was grown and prepared as described above (Materials and Methods 2a). The *Burkholderia* and *Ralstonia* isolates were grown in separate tubes, washed and added to the rest of the SynCom to a final OD of 0.001 (the calculated OD of each individual strain in a 185-Member SynCom at an OD of 0.2), to form the following four mixtures: (1) Full community – all *Burkholderia* and *Ralstonia* isolates added to the SynCom; (2) *Burkholderia* drop-out – only *Ralstonia* isolates added to the SynCom; (3) *Ralstonia* drop-out – only *Burkholderia* isolates added to the SynCom; (4) Uninoculated plants – no SynCom. The experiment had three Pi conditions: low Pi (50 µM Pi), high Pi (1000 µM Pi) and low→high Pi. 12 days post-inoculation the low Pi and high Pi samples were harvested, and the low→high plants were transferred from 50 µM Pi plates to 1000 µM Pi plates for an additional 3 days. The experiment was performed twice and each rep consisted of six plates per SynCom mixture and Pi treatment, amounting to 72 samples. Upon harvest, shoot Pi accumulation was measured using the Ames method (Materials and Methods 1g).

#### c. In vitro plant growth conditions

All seeds were surface-sterilized with 70% bleach, 0.2% Tween-20 for 8 min, and rinsed three times with sterile distilled water to eliminate any seed-borne microbes on the seed surface. Seeds were stratified at 4 °C in the dark for two days. Plants were germinated on vertical square 10 × 10 cm agar plates with Johnson medium (JM; [4]) containing 0.5% sucrose and 1000 µM Pi, for 7 days. Then, 10 plants were transferred to each vertical agar plate with amended JM lacking sucrose at one of the following experimental Pi concentrations: 0, 10, 30, 50, 100 or 1000 µM Pi. The SynCom was spread on the agar prior to transferring plants. Each experiment included unplanted agar plates with SynCom for each media type (designated NP) and uninoculated plates with plants for each media type (designated NB). Plants were placed in randomized order in growth chambers and grown under a 16-h dark/8-h light regime at 21°C day/18°C night for 12 days.

#### d. Sample harvest

Twelve days post-transferring, plates were imaged using a document scanner. For DNA extraction; roots, shoots and agar were harvested separately, pooling 6 plants for each sample. Roots and shoots were placed in 2.0 ml Eppendorf tubes with three sterile glass beads. Samples were washed three times with sterile distilled water to remove agar particles and weakly associated microbes. Tubes were stored at −80 °C until processing. For RNA, samples were collected from a separate set of two independent experiments, using the same SynCom and conditions as above, but with just two Pi concentrations: 1000 µM Pi (high) and 50 µM Pi (low). Four seedlings were harvested from each sample and samples were flash frozen and stored at −80 °C until processing.

#### e. DNA extraction

Root and shoot samples were lyophilized for 48 hours using a Freezone 6 freeze dry system (Labconco) and pulverized using a tissue homogenizer (MPBio). Agar from each plate was stored in a 30 ml syringe with a square of sterilized Miracloth (Millipore) at the bottom and kept at −20 °C for a week. Syringes were then thawed at room temperature and samples were squeezed gently into 50 ml tubes. Samples were centrifuged at maximum speed for 20 minutes and most of the supernatant was discarded. The remaining 1-2 ml of supernatant containing the pellet was transferred into clean microfuge tubes. Samples were centrifuged again, supernatant was removed, and pellets were stored at −80 °C until DNA extraction.

DNA extractions were carried out on ground root and shoot tissue and agar pellets using 96-well-format MoBio PowerSoil Kit (MOBIO Laboratories; Qiagen) following the manufacturer’s instruction. Sample position in the DNA extraction plates was randomized, and this randomized distribution was maintained throughout library preparation and sequencing.

#### f. RNA extraction

RNA was extracted from *A*. *thaliana* seedlings following [56]. Frozen seedlings were ground in liquid nitrogen, then homogenized in a buffer containing 400 μl of Z6-buffer; 8 M guanidinium-HCl, 20 mM MES, 20 mM EDTA at pH 7.0. 400 μL phenol:chloroform:isoamylalcohol, 25:24:1 was added, and samples were vortexed and centrifuged (20,000 g, 10 minutes) for phase separation. The aqueous phase was transferred to a new 1.5 ml tube and 0.05 volumes of 1 N acetic acid and 0.7 volumes 96% ethanol were added. The RNA was precipitated at −20 °C overnight. Following centrifugation (20,000 g, 10 minutes, 4 °C), the pellet was washed with 200 μl sodium acetate (pH 5.2) and 70% ethanol. The RNA was dried and dissolved in 30 μL of ultrapure water and stored at −80 °C until use.

#### g. Quantification of plant phenotypes

The Ames method [55] was used to determine the phosphate concentration in the shoots of plants grown on different Pi regimens and treatments. Primary root length elongation was measured using ImageJ [57] and for shoot area and total root network measurement, WinRhizo software (Regens Instruments Inc.), was used.

### 3. DNA and RNA sequencing

#### a. Bacterial 16S sequencing

We amplified the V3-V4 regions of the bacterial 16S rRNA gene using primers 338F (5′-ACTCCTACGGGAG GCAGCA-3′) and 806R (5′-GGACTACHVGGGTWTCTAAT-3′). Two barcodes and 6 frameshifts were added to the 5’ end of 338F and 6 frameshifts were added to the 806R primers, based on the protocol in [58]. Each PCR reaction was performed in triplicate, and included a unique mixture of three frameshifted primer combinations for each plate. PCR conditions were as follows: 5 μl Kapa Enhancer (Kapa Biosystems), 5 μl Kapa Buffer A, 1.25 μl of 5 μM 338F, 1.25 μl of 5 μM 806R, 0.375 μl mixed rRNA gene-blocking peptide nucleic acids (PNAs; 1:1 mix of 100 μM plastid PNA and 100 μM mitochondrial PNA; PNA Bio), 0.5 μl Kapa dNTPs, 0.2 μl Kapa Robust Taq, 8 μl dH2O, 5 μl DNA; temperature cycling: 95 °C for 60 s, 24 cycles of 95 °C for 15 s, 78 °C (PNA) for 10 s, 50 °C for 30 s, 72 °C for 30 s, 4 °C until use. Following PCR cleanup, the PCR product was indexed using 96 indexed 806R primers with the same reaction mix as above, and 9 cycles of the cycling conditions described in [29]. PCR products were purified using AMPure XP magnetic beads (Beckman Coulter) and quantified with a Qubit 2.0 fluorometer (Invitrogen). Amplicons were pooled in equal amounts and then diluted to 10 pM for sequencing. Sequencing was performed on an Illumina MiSeq instrument using a 600-cycle V3 chemistry kit. The raw data for the natural soil experiment is available in the NCBI SRA Sequence Read Archive (accession XXXXXXX). The raw data for the SynCom experiment is available in the NCBI SRA Sequence Read Archive (accession XXXXXXX).

#### b. Fungal/Oomycete ITS sequencing

We amplified the ITS1 region using primers ITS1-F (5′-CTTGGTCATTTAGAGGAAGTAA-3′; [59]) and ITS2 (5′-GCTGCGTTCTTCATCGATGC-3′; [60]). Samples were diluted to concentrations of 3.5 ng µl^−1^ of DNA with nuclease-free water for the first PCR reaction to amplify the ITS1 region. Reactions were prepared in triplicate in 25 µl volumes consisting of 10 ng of DNA template, 1× incomplete buffer, 0.3% bovine serum albumin, 2 mM MgCl_2_, 200 µM dNTPs, 300 nM of each primer and 2 U of DFS-Taq DNA polymerase (Bioron, Ludwigshafen, Germany); temperature cycling: 2 min at 94 °C, 25 cycles: 30 s at 94 °C, 30 s at 55 °C, and 30 s at 72 °C; and termination: 10 min at 72 °C. PCR products were cleaned using an enzymatic cleanup (24.44 µl: 20 µl of template, 20 U of exonuclease I, 5 U of Antarctic phosphatase, 1× Antarctic phosphatase buffer; New England Biolabs, Frankfurt, Germany); incubation conditions: (30 min at 37 °C, 15 min at 85 °C; centrifuge 10 min at 4,000 rpm). A second PCR was then performed (2 min at 94 °C; 10 cycles: 30 s at 94 °C, 30 s at 55 °C, and 30 s at 72 °C; and termination: 10 min at 72 °C), in triplicate using 3 µl of cleaned PCR product and sample-specific barcoded primers (5′-AATGATACGGCGACCACCGAGATCTACACTCACGCGCAGG-ITS1F-3′; 5′-CAAGCAGAAGACGGCATACGAGAT-BARCODE(12-NT)-CGTACTGTGGAGA-ITS2-3′). PCR reactions were purified using with Agencourt AMPure XP purification kit (Beckman Coulter, Krefeld, Germany). Amplicons were pooled in equal amounts and then diluted to 10 pM for sequencing. Sequencing was performed on an Illumina MiSeq instrument using a 600-cycle V3 chemistry kit. The raw data is available in the NCBI SRA Sequence Read Archive (Project Number PRJNA531340).

#### c. Plant RNA sequencing

Illumina-based mRNA-Seq libraries were prepared from 1 μg RNA following [51]. mRNA was purified from total RNA using Sera-mag oligo(dT) magnetic beads (GE Healthcare Life Sciences) and then fragmented in the presence of divalent cations (Mg^2+^) at 94 °C for 6 minutes. The resulting fragmented mRNA was used for first-strand cDNA synthesis using random hexamers and reverse transcriptase, followed by second-strand cDNA synthesis using DNA Polymerase I and RNAseH. Double-stranded cDNA was end-repaired using T4 DNA polymerase, T4 polynucleotide kinase, and Klenow polymerase. The DNA fragments were then adenylated using Klenow exo-polymerase to allow the ligation of Illumina Truseq HT adapters (D501– D508 and D701–D712). All enzymes were purchased from Enzymatics. Following library preparation, quality control and quantification were performed using a 2100 Bioanalyzer instrument (Agilent) and the Quant-iT PicoGreen dsDNA Reagent (Invitrogen), respectively. Libraries were sequenced using Illumina HiSeq4000 sequencers to generate 50-bp single-end reads.

### 4. Data processing and Statistical analyses

#### a. Quantification of plant phenotypes – soil experiment

To measure correlation between all measured plant phenotypes (shoot Pi, shoot weight, shoot size) we applied hierarchical clustering based on the all vs all pairwise correlation coefficients between all the phenotypes measured. We used the R package corrplot v.0.84 [61] to visualize correlations. To compare shoot Pi accumulation, we performed a paired t-test between low P and P-supplemented low P (low+P) samples, within each plant genotype independently (α < 0.05).

#### b. Amplicon sequence data processing – soil experiments

Bacterial sequencing data was processed with MT-Toolbox [62]. Usable read output from MT-Toolbox (that is, reads with 100% correct primer and primer sequences that successfully merged with their pair) were quality filtered using Sickle [63] by not allowing any window with a Q-score under 20. After quality filtering, samples with < 3000 reads, amounting to 51 samples, all soil samples, were discarded. The resulting sequences were collapsed into amplicon sequence variants (ASVs) using the R package DADA2 v1.8.1 [64]. Taxonomic assignment of each ASV was performed using the naïve Bayes kmer method implemented in the DADA2 package using the Silva 132 database as training reference [64].

Fungal sequencing data was processed as previously described [41]. Briefly, a combination of QIIME [65] and USEARCH [66] pipelines were used to cluster the fungal reads into 97% OTUs. Filtering of non-fungal OTUs was performed by aligning each representative against a dedicated ITS database. Finally, taxonomic assignment of each OTU was performed using the WarCup fungal ITS training set (2016) [67].

The resulting bacterial and fungal count tables were deposited at https://github.com/isaisg/hallepi

#### c. Community analyses – soil experiments

The resulting bacterial and fungal count tables were processed and analyzed with functions from the ohchibi package [68]. Both tables were rarefied to 3000 reads per sample. An alpha diversity metric (Shannon diversity) was calculated using the diversity function from the vegan package v2.5-3 [69]. We used ANOVA to test for differences in Shannon Diversity indices between groups. Tukey’s HSD post-hoc tests here and elsewhere were performed using the cld function from the emmeans R package [70]. Beta diversity analyses (Principal coordinate analysis, and canonical analysis of principal coordinates) were based on Bray-Curtis dissimilarity calculated from the rarefied abundance tables. We utilized the capscale function from the vegan R package v.2.5-3 [69] to compute a constrained analysis of principal coordinates (CAP). To analyze the full dataset (all fraction, all genotypes all phosphorus treatments), we constrained by fraction, plant genotype and phosphorus fertilization treatment, while conditioning for the plot effect. We performed the Genotype: phosphorus interaction analysis over each fraction independently, constraining for the plant genotype and phosphorus fertilization treatment while conditioning for the plot effect. In addition to CAP, we performed Permutational Multivariate Analysis of Variance (PERMANOVA) over the two datasets described above using the adonis function from the vegan package v2.5-3 [69]. Finally, we used the function chibi.permanova from the ohchibi package to plot the R^2^ values for each significant term in the PERMANOVA model tested.

The relative abundance of bacterial phyla and fungal taxa were depicted using the stacked bar representation encoded in the function chibi.phylogram from the ohchibi package.

We used the R package DESeq2 v1.22.1 [71] to compute the enrichment profiles for both bacterial ASVs and fungal OTUs. For the full dataset model, we estimated main effects for each variable tested (Fraction, Plant Genotype, and Phosphorus fertilization) using the following design:

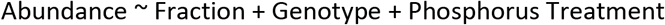

We delimited ASV/OTU fraction enrichments using the following contrasts: Soil vs Root, Soil vs Shoot and Root vs Shoot. An ASV/OTU was considered statistically significant if it had q-value < 0.1.

We implemented a second statistical model in order to identify ASVs and OTUs that exhibited statistically significant differential abundances depending on genotype. For this analysis we utilized only root-derived low P and P-supplemented low P (low+P) treatments. We utilized a group design framework to facilitate the construction of specific contrasts. In the group variable we created, we merged the genotype and phosphate levels per sample (e.g. Col-0_lowP, *phf1*_low+P or *phr1 phl1*_lowP). We controlled the paired structure of our design by adding a plot variable, resulting in the following model design:

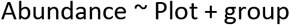

We delimited 6 sets (S1, S2, S3, S4, S5, S6) of statistically significant (q-value < 0.1) ASVs/OTUs from our model using the following contrasts:

*S1* = {Samples from Col-0, higher abundance in low treatment in comparison to low+P treatment}

*S2* = {Samples from *phf1*, higher abundance in low treatment in comparison to low+P treatment}

*S3* = {Samples from *phr1 phl1*, higher abundance in low treatment in comparison to low+P treatment}

*S4* = {Samples from *Col-0*, higher abundance in low+P treatment in comparison to low treatment}

*S5* = {Samples from *phf1*, higher abundance in low+P treatment in comparison to low treatment}

*S6* = {Samples from *phr1 phl1*, higher abundance in low+P treatment in comparison to low treatment}

The six sets described above were used to populate Figures 3c-f.

The interactive visualization of the enrichment profiles was performed by converting the taxonomic assignment of each ASV/OTU into a cladogram with equidistant branch lengths using R. We used the interactive tree of life interface (iTOL) [72] to visualize this tree jointly with metadata files derived from the output of the statistical models described above. The cladograms for both bacteria and fungi can be downloaded from the links described above or via the iTOL user hallepi.

In order to compare beta diversity patterns across samples, we only used samples coming from pots where sequence data from all three fractions (soil root and shoot) passed quality filtering. Then, for each fraction we estimated a distance structure between samples inside that fraction using the Bray Curtis dissimilarity metric. Finally, we computed Mantel [73] correlations between pairs of distance objects (e.g. samples from Root or samples from Shoot) using the vegan package v2.5-3 [69] implementation of the Mantel test.

All scripts and datasets required to reproduce the soil experiment analyses are deposited in the following GitHub repository: https://github.com/isaisg/hallepi/.

#### d. Amplicon sequence data processing – SynCom experiments

SynCom sequencing data were processed with MT-Toolbox [62]. Usable read output from MT-Toolbox (that is, reads with 100% correct primer and primer sequences that successfully merged with their pair) were quality filtered using Sickle [63] by not allowing any window with Q-score under 20. The resulting sequences were globally aligned to a reference set of 16S rRNA gene sequences extracted from genome assemblies of SynCom member strains. For strains that did not have an intact 16S rRNA gene sequence in their assembly, we generated the 16S rRNA gene using Sanger sequencing. The reference database also included sequences from known bacterial contaminants and Arabidopsis organellar 16S sequences (S12 table). Sequence alignment was performed with USEARCH v7.1090 [66] with the option ‘usearch_global’ at a 98% identity threshold. On average, 85% of sequences matched an expected isolate. Our 185 isolates could not all be distinguished from each other based on the V3-V4 sequence and were thus classified into 97 unique sequences (USeqs). A USeq encompasses a set of identical (clustered at 100%) V3-V4 sequences coming from a single or multiple isolates.

Sequence mapping results were used to produce an isolate abundance table. The remaining unmapped sequences were clustered into Operational Taxonomic Units (OTUs) using UPARSE [74] implemented with USEARCH v7.1090, at 97% identity. Representative OTU sequences were taxonomically annotated with the RDP classifier [75] trained on the Greengenes database [76] (4 February 2011). Matches to Arabidopsis organelles were discarded. The vast majority of the remaining unassigned OTUs belonged to the same families as isolates in the SynCom. We combined the assigned Useq and unassigned OTU count tables into a single table.

The resulting count table was processed and analyzed with functions from the ohchibi package. Samples were rarefied to 1000 reads per sample. An alpha diversity metric (Shannon diversity) was calculated using the diversity function from the vegan package v2.5-3 [69]. We used ANOVA to test for differences in alpha diversity between groups. Beta diversity analyses (Principal coordinate analysis, and canonical analysis of principal coordinates) were based on were based on Bray-Curtis dissimilarity calculated from the rarefied abundance tables. We used the capscale function from the vegan R package v.2.5-3 [69] to compute the canonical analysis of principal coordinates (CAP). To analyze the full dataset (all fraction, all phosphate treatments), we constrained by fraction and phosphate concentration while conditioning for the replicate effect. We performed the Fraction:Phosphate interaction analysis within each fraction independently, constraining for the phosphate concentration while conditioning for the rep effect. In addition to CAP, we performed Permutational Multivariate Analysis of Variance (PERMANOVA) analysis over the two datasets described above using the adonis function from the vegan package v2.5-3 [69]. Finally, we used the function chibi.permanova from the ohchibi package to plot the R^2^ values for each significant term in the PERMANOVA model tested.

We visualized the relative abundance of the bacterial phyla present in the SynCom using the stacked bar representation encoded in the chibi.phylogram from the ohchibi package.

We used the package DESeq2 v1.22.1 [71] to compute the enrichment profiles for USeqs and OTUs present in the count table. For the full dataset model, we estimated main effects for each variable tested (fraction and phosphate concentration) using the following model specification:

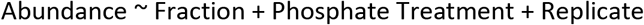

We calculated the USeqs/OTUs fraction enrichments using the following contrasts: Agar vs Root, Agar vs Shoot and Root vs Shoot. A USeq/OTU was considered statistically significant if it had q-value < 0.1. In order to populate the heatmaps shown in Figure 5C, we grouped the Fraction and Phosphate treatment variable into a new group variable that allowed us to fit the following model:

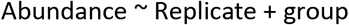

We used the fitted model to estimate the fraction effect inside each particular phosphate level (e.g. Root vs Agar at 0Pi, or Shoot vs Agar at 1000Pi).

Additionally, we utilized a third model for the identification of USeqs/OTUs that exhibited a significant Fraction:Phosphate interaction between the planted agar samples and the plant fractions (Root and Shoot). Based on the beta diversity and alpha diversity results, we only used samples that were treated with 0, 10, 100 and 1000 µM of phosphate. We grouped the samples into two categories based on their phosphate concentration, low (0 µM and 10 µM) and high (100 µM and 1000 µM). Then we used the following model specification to derive the desired interaction effect:

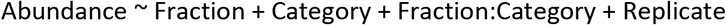

Finally, we subset USeqs that exhibited a significant interaction (Fraction:Category, q-value < 0.1) in the following two contrasts (Planted Agar vs Root) and (Planted Agar vs Shoot).

In order to compare beta diversity patterns across samples, we only used samples coming from pots where sequence data from all three fractions (soil root and shoot) passed quality filtering. Then, for each fraction we estimated a distance structure between samples inside that fraction using the Bray Curtis dissimilarity metric. Finally, we computed Mantel [73] correlations between pairs of distance objects (e.g. samples from Root or samples from Shoot) using the vegan package v2.5-3 [69] implementation of the Mantel test.

For the drop-out experiment, we ran an ANOVA model inside each of the phosphate treatments tested (50 µM Pi, 1000 µM Pi and 50→1000 µM Pi). We visualized the results of the ANOVA models using the compact letter display encoded in the CLD function from the emmeans package.

All scripts necessary to reproduce the synthetic community analyses are deposited in the following GitHub repository: https://github.com/isaisg/hallepi.

#### e. Phylogenetic inference of the SynCom Isolates

To build the phylogenetic tree of the SynCom isolates, we utilized the super matrix approach previously described in [47]. Briefly, we scanned 120 previously defined marker genes across the 185 isolate genomes from the SynCom utilizing the hmmsearch tool from the hmmer v3.1b2 [77]. Then, we selected 47 markers that were present as single copy genes in 100% of our isolates. Next, we aligned each individual marker using MAFFT [78] and filtered low quality columns in the alignment using trimAl [79]. Afterwards, we concatenated all filtered alignments into a super alignment. Finally FastTree v2.1 [80] was used to infer the phylogeny utilizing the WAG model of evolution.

We utilized the inferred phylogeny along with the fraction fold change results of the main effect model to compute the phylogenetic signal (Pagel’s λ) [81] for each contrast (Planted Agar vs Root, Planted Agar vs Shoot and Root vs Shoot) along each concentration of the phosphate gradient. The function phylosig from the R package phytools [82] was used to test for significance of the phylogenetic signal measured.

Multiple panel figures were constructed using the egg R package [83].

#### f. RNA-Seq read processing

Initial quality assessment of the Illumina RNA-seq reads was performed using FastQC v0.11.7 [84]. Trimmomatic v0.36 [85] was used to identify and discard reads containing the Illumina adaptor sequence. The resulting high-quality reads were then mapped against the TAIR10 [86] Arabidopsis reference genome using HISAT2 v2.1.0 [87]with default parameters. The featureCounts function from the Subread package [88] was then used to count reads that mapped to each one of the 27,206 nuclear protein-coding genes. Evaluation of the results of each step of the analysis was done with MultiQC v1.1 [89]. Raw sequencing data and read counts are available at the NCBI Gene Expression Omnibus accession number GSE129396.

#### g. RNA-Seq statistical analysis – soil experiment

To measure the transcriptional response to Pi limitation in soil, we used the package DESeq2 v.1.22.1 [71] to define differentially expressed genes (DEGs) using the raw count table described above (Materials and Methods 4f). We used only samples from low P and P-supplemented low P (low+P) treatments along the three genotypes tested (Col-0, *phf1* and *phr1 phl1*). We combined the Genotype and Phosphorus Treatment variables into a new group variable (e.g. Col-0_lowP or *phf1*_low+P). Because we were interested in identifying DEGs among any pair of levels (6 levels) of the group variable (e.g. Col-0_lowP vs Col-0_low+P) we performed a likelihood ratio test (LRT) between a model containing the group variable and a reduced model containing only the intercept. Next, we defined DEGs as genes that had a q-value < 0.1.

For visualization purposes, we applied a variance stabilizing transformation to the raw count gene matrix. We then standardized (z-score) each gene along the samples. We subset DEGs from this standardized matrix and for each gene calculated the mean z-score expression value in a particular level of the group variable (e.g. Col-0_lowP); this resulted in a matrix of DEGs across the six levels in our design. Next, we created a dendrogram of DEGs by applying hierarchical clustering (method ward.D2, hclust R-base [90]) to a distance object based on the correlation (dissimilarity) of the expression profiles of the genes across the six levels in our design. Finally, we delimited the cluster of DEGs by cutting the output dendogram into five groups using the R-base cutree function [90]. Gene ontology enrichment was performed for each cluster of DEGs using the R package clusterProfiler [91].

For the PSR marker gene analysis we downloaded the ID of 193 genes defined in [4]. Then, we subset these genes from our standardized matrix and computed for each gene the mean z-score expression value in a particular level of the group variable. Finally, we visualized the average expression of this PSR regulon across our groups of interest utilizing the function chibi.boxplot from the ohchibi package.

All scripts necessary to reproduce the RNA-Seq analyses are deposited in the following GitHub repository: https://github.com/isaisg/hallepi.

#### h. RNA-Seq statistical analysis – SynCom experiment

To measure the transcriptional response to Pi limitation in the SynCom microcosm, we used the package DESeq2 v.1.22.1 [71] to define differentially expressed genes (DEGs) using the raw count gene table. We combined the Bacteria (No bacteria, Full SynCom) and Phosphorus Treatment variables into a new group variable (e.g. NB_50Pi or Full_1000Pi). Afterwards we fitted the following model to our gene matrix:

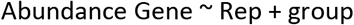

Finally, utilizing the model fitted, we contrasted the phosphate treatment inside each level of the Bacteria variable (e.g. NB_1000Pi vs NB_50Pi). Any gene with q-value < 0.1 was defined as differentially expressed.

For the PSR marker gene analysis we downloaded the ID of 193 genes defined in [4]. Then, we subset these genes from our standardized matrix and computed for each gene the mean z-score expression value in a particular level of the group variable. Finally, we visualized the average expression of the PSR regulon across our groups of interest utilizing the function chibi.boxplot from the ohchibi package.

All scripts necessary to reproduce the RNA-Seq analyses are deposited in the following GitHub repository: https://github.com/isaisg/hallepi

## Acknowledgments

We thank Dr. Wolfgang Gans and Prof. Edgar Peiter (Martin Luther University Halle-Wittenberg, Halle, Germany) for generously providing us with access to Halle soils; Prof. Paul Schulze-Lefert (Max Planck Institute for Plant Breeding Research, Cologne, Germany) for generously providing greenhouse and lab space; Stratton Barth, Julia Shen, Ellie Wilson, May Priegel and Dilan Chudasma for technical assistance throughout the project; the Dangl lab microbiome group for useful discussions; Dr. Javier Paz-Ares, Dr. Antonio Leyva (National Centre for Biotechnology, Madrid, Spain) and Dr. Connor Fitzpatrick, for critical comments on the manuscript. This work was supported by NSF INSPIRE grant IOS-1343020 and DOE-USDA Feedstock Award DE-SC001043 to J.L.D. J.L.D is an Investigator of the Howard Hughes Medical Institute, supported by the HHMI. P.J.P.T.L was supported by the Pew Latin American Fellows Program in the 746 Biomedical Sciences (grant number 00026198). O.M.F was supported by NIH NRSA Fellowship F32-GM117758.

---

**S1 Figure.**
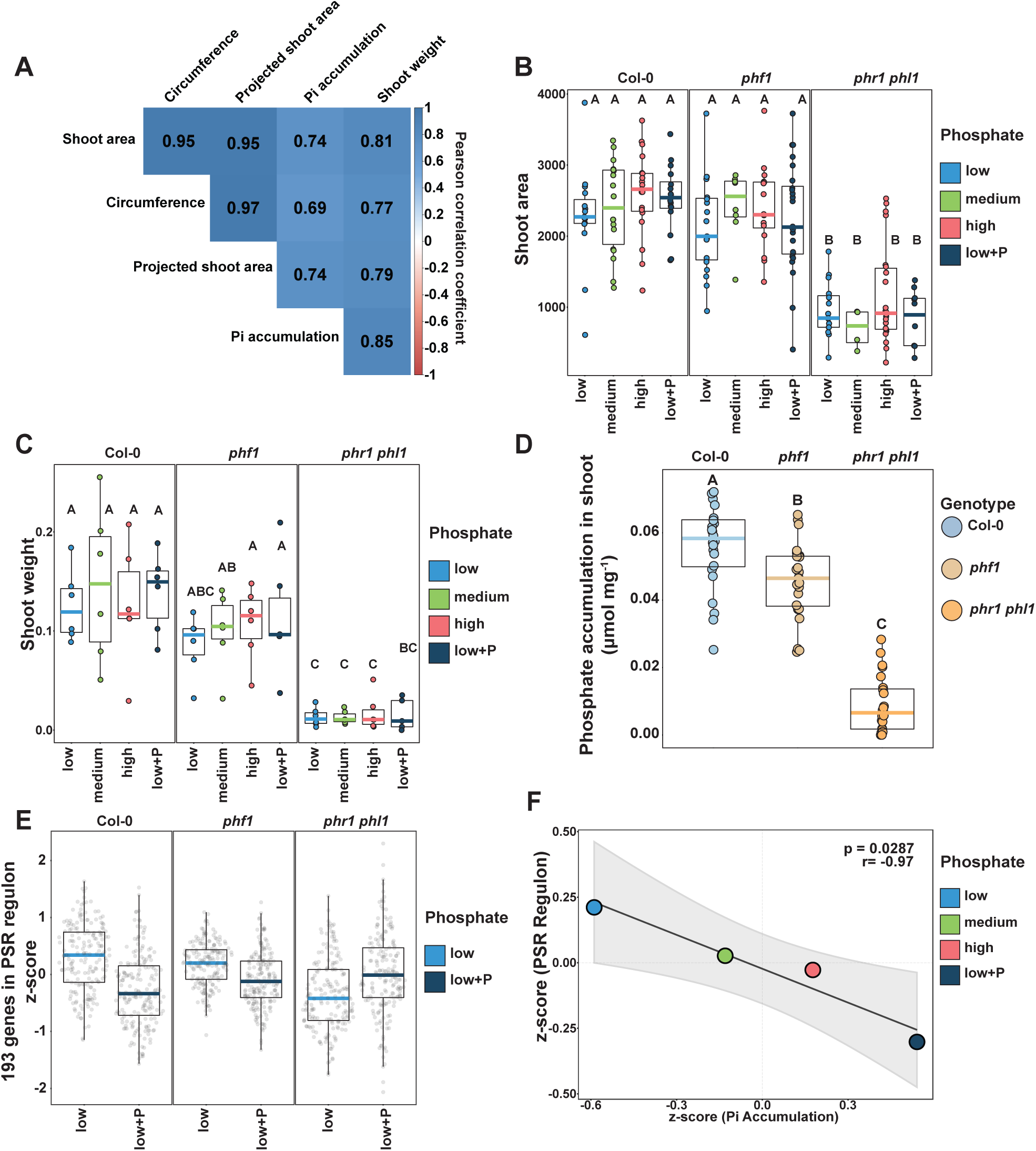
Phosphate starvation response in soil. **(A)** Heatmap showing the all vs all pairwise Pearson correlation coefficient calculated between the quantified phenotypes associated with the phosphate starvation response: shoot area, shoot fresh weight and shoot free Pi accumulation. **(B)** Boxplot showing the distribution of the shoot area measured across the phosphorus gradient within each of the three genotypes. Lefters represent the results of the post hoc test. **(C)** Boxplot showing the distribution of shoot fresh weight measured across the phosphorus gradient within each of the three genotypes. Lefters illustrate the results of the post hoc test. **(D)** Boxplot showing the shoot Pi accumulation across the three genotypes. Lefters represent the results of the post hoc test. **(E)** Boxplots displaying the average expression of 193 PSR marker genes across the low and low+P samples in each of the three genotypes tested. **(F)** Scatterplot showing the relationship between the standardized average phosphate accumulation in leaves (x-axis) and the average standardized expression of 193 PSR marker genes (y-axis). The p-value and R value were calculated according to Pearson’s product moment correlation coefficient.

**S2 Figure.**
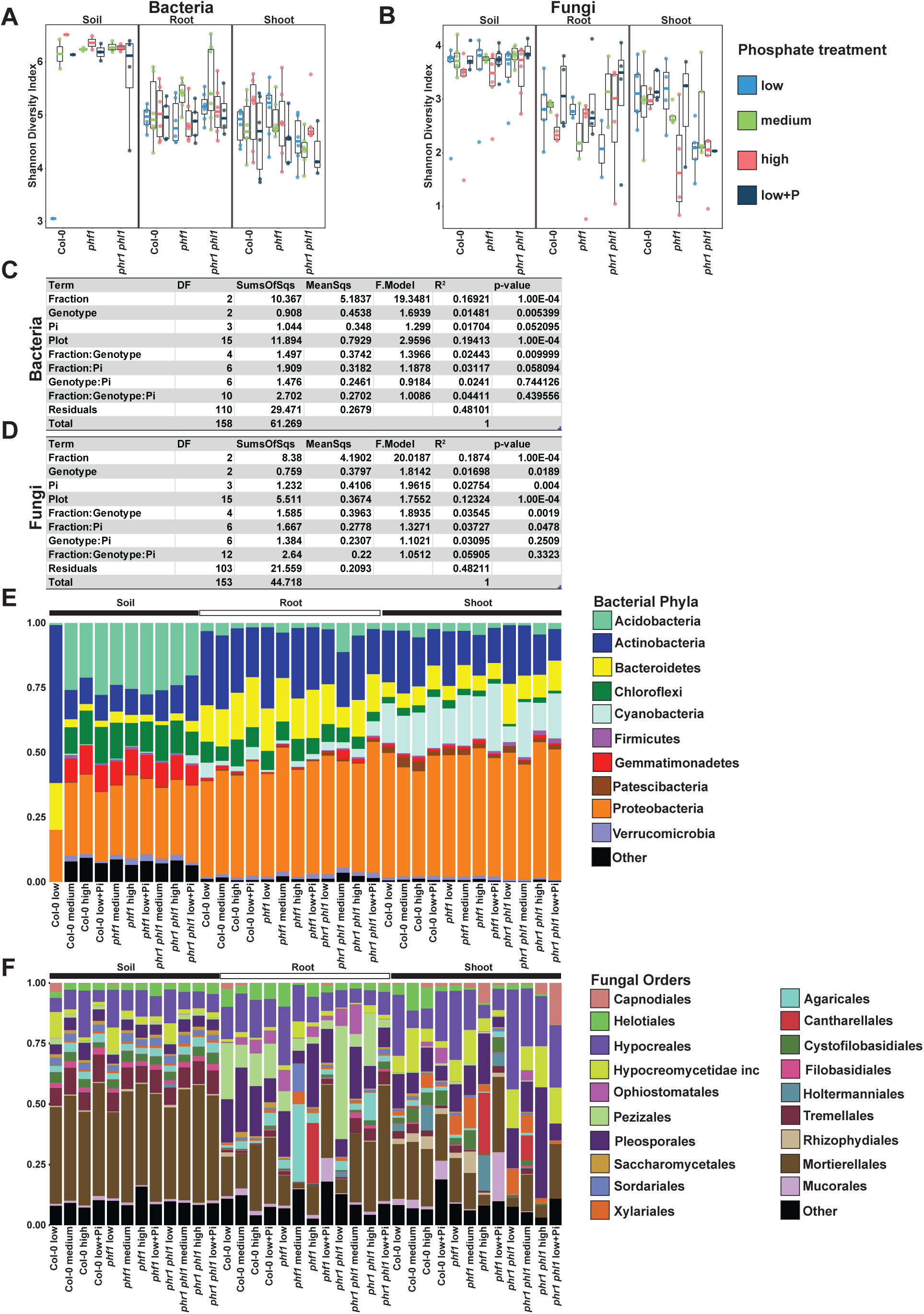
Characterization of the soil and plant microbiota in soils exposed to different level of phophorus fertilization. **(A,B)** Boxplots showing the distribution of the alpha diversity (Shannon diversity index) across all levels of phosphorus in the soil for bacteria **(A)** and fungi **(B)**. **(C,D)** PERMANOVA results in which the effect of the three variables (Fraction, Genotype and Soil P) and their interaction on the assembly of the bacterial **(C)** and fungal **(D)** communities were tested. **(E)** Relative abundance profiles of the main bacterial phyla in the three variables (Fraction, Genotype and Soil P) across all levels of P in the soils. **(F)** Relative abundance profiles of the main fungal orders in the three variables (Fraction, Genotype and Soil P) across all the levels of P in the soils.

**S3 Figure.**
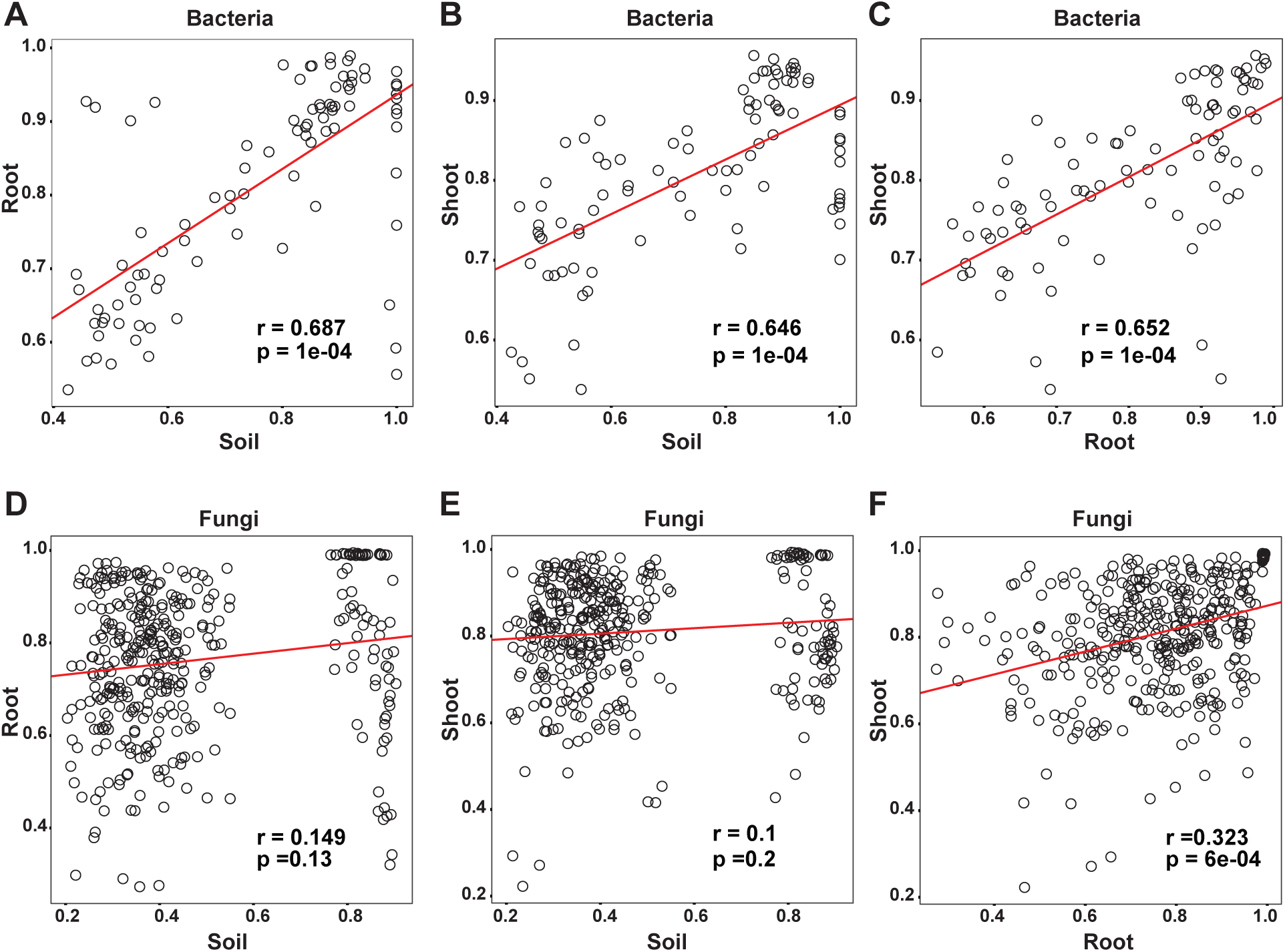
Bacterial but not fungal plant microbiota composition is strongly dependent on soil inoculum. **(A,B,C)** Correlation plots between Bray-Curtis distance matrices calculated for bacteria within soil treatments, root and shoot fractions. The R and p-values were calculated using Mantel tests. **(A)** Correlation plot of soil vs root. **(B)** Correlation plot of soil vs shoot. **(C)** Correlation plot of root vs shoot. **(D,E,F)** Correlation plots between Bray-Curtis distance matrices calculated for fungi within soil treatments, root and shoot fractions. The R and p-values were calculated using Mantel tests. **(D)** Correlation plot of soil vs root. **(E)** Correlation plot of soil vs shoot. **(F)** Correlation plot of root vs shoot.

**S4 Figure.**
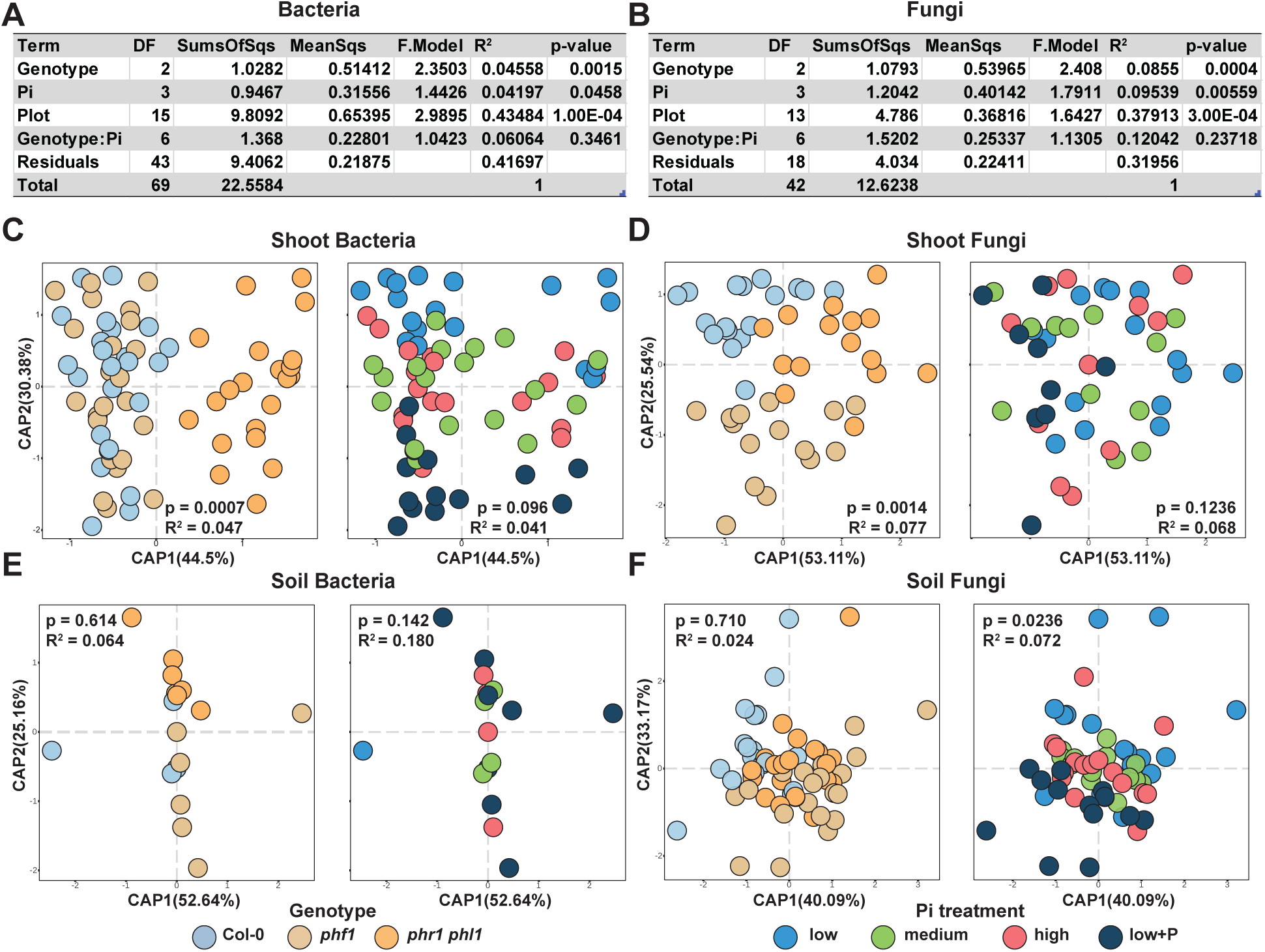
Plant genotypes and soil phosphorus concentrations influence the composition of the plant microbiota. **(A,B)** PERMANOVA results showing the influence of the plant genotype and soil P concentration and their interaction on the assembly of the root **(A)** bacterial and **(B)** fungal communities. **(C,D)** Canonical analysis of principal coordinates showing the effect of plant genotype and P content in the soil over the shoot **(C)** bacterial and **(D)** fungal communities (Materials and methods 4b). The p-value and R2 values in each plot are derived from a PERMANOVA model and correspond to the genotype and soil P term respectively. **(E,F)** Canonical analysis of principal coordinates showing the influence of genotype and P on the soil **(E)** bacterial and **(F)** fungal communities (Materials and methods 4b). The p-value and R2 values in each plot are derived from a PERMANOVA model and correspond to the genotype and soil P term respectively.

**S5 Fig.**
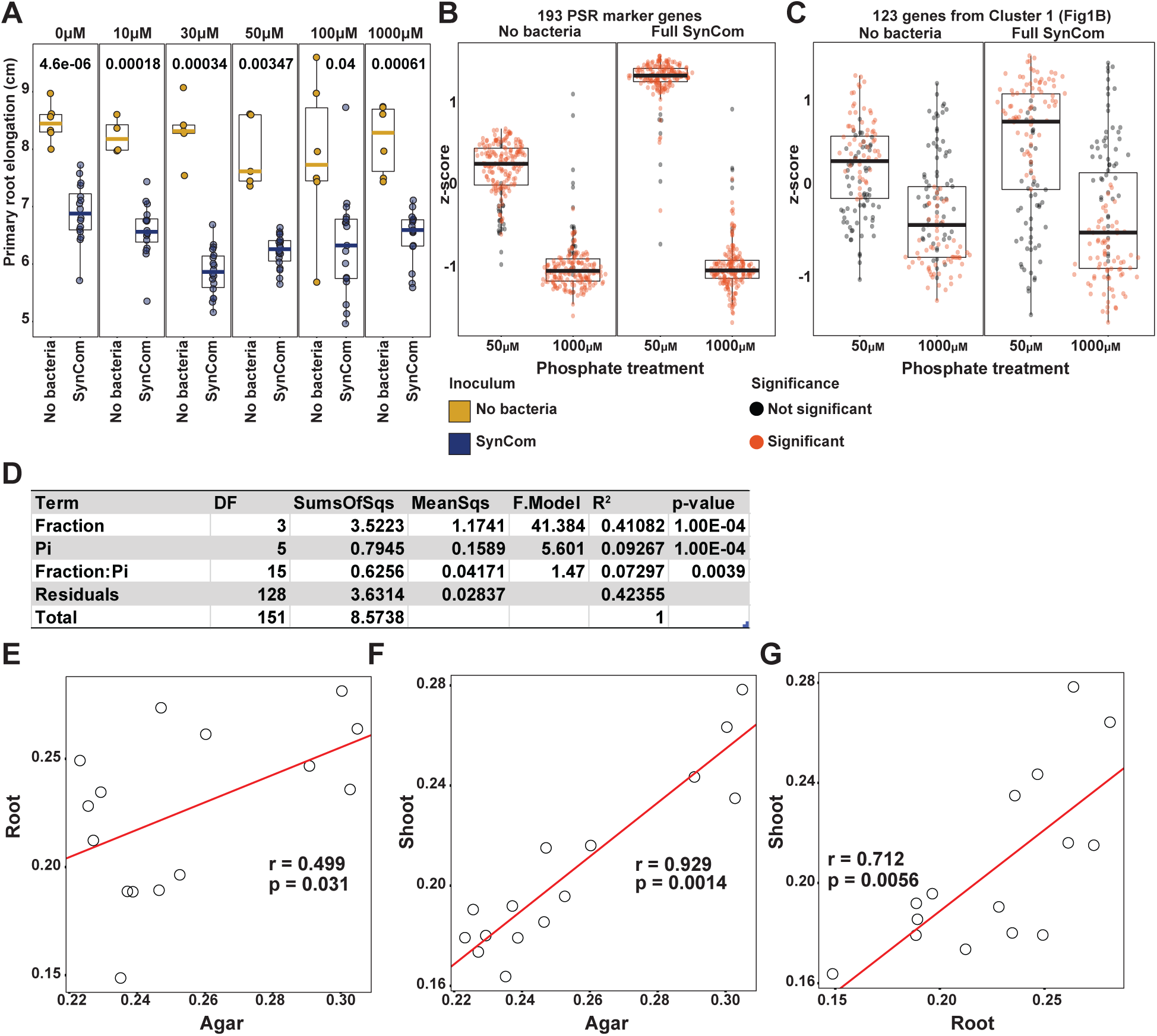
A bacterial synthetic community modifies the plant phosphate starvation response. **(A)** Boxplots displaying the primary root elongation of plants grown in a gradient of Pi concentrations in sterile conditions or with the SynCom (Materials and methods 2g). A t-test was used for each Pi treatment to estimate differences between SynCom-treated and uninoculated plants. **(B)** Average expression of the 193 PSR markers genes in low (50 µM) and high (1000 µM) Pi conditions within Syncom-treated and uninoculated plants. (Materials and methods 4g). (**C)** Average expression of the 123 genes from Cluster 1 (Figure 1B) in low (50 µM) and high (1000 µM) Pi conditions within Syncom-treated and uninoculated plants. **(D)** PERMANOVA model results showing the influence of the two variables (Fraction and Pi concentration) and their interaction on the assembly of the bacterial community in the plant. **(E,F,G)** Correlation plots between Bray-Curtis distance matrices calculated for bacterial profile within agar, root and shoot fractions. The R and p-values were calculated using mantel tests. **(E)** Correlation plot of agar vs root. **(F)** Correlation plot of agar vs shoot. **(G)** Correlation plot of root vs shoot.

**S6 Figure.**
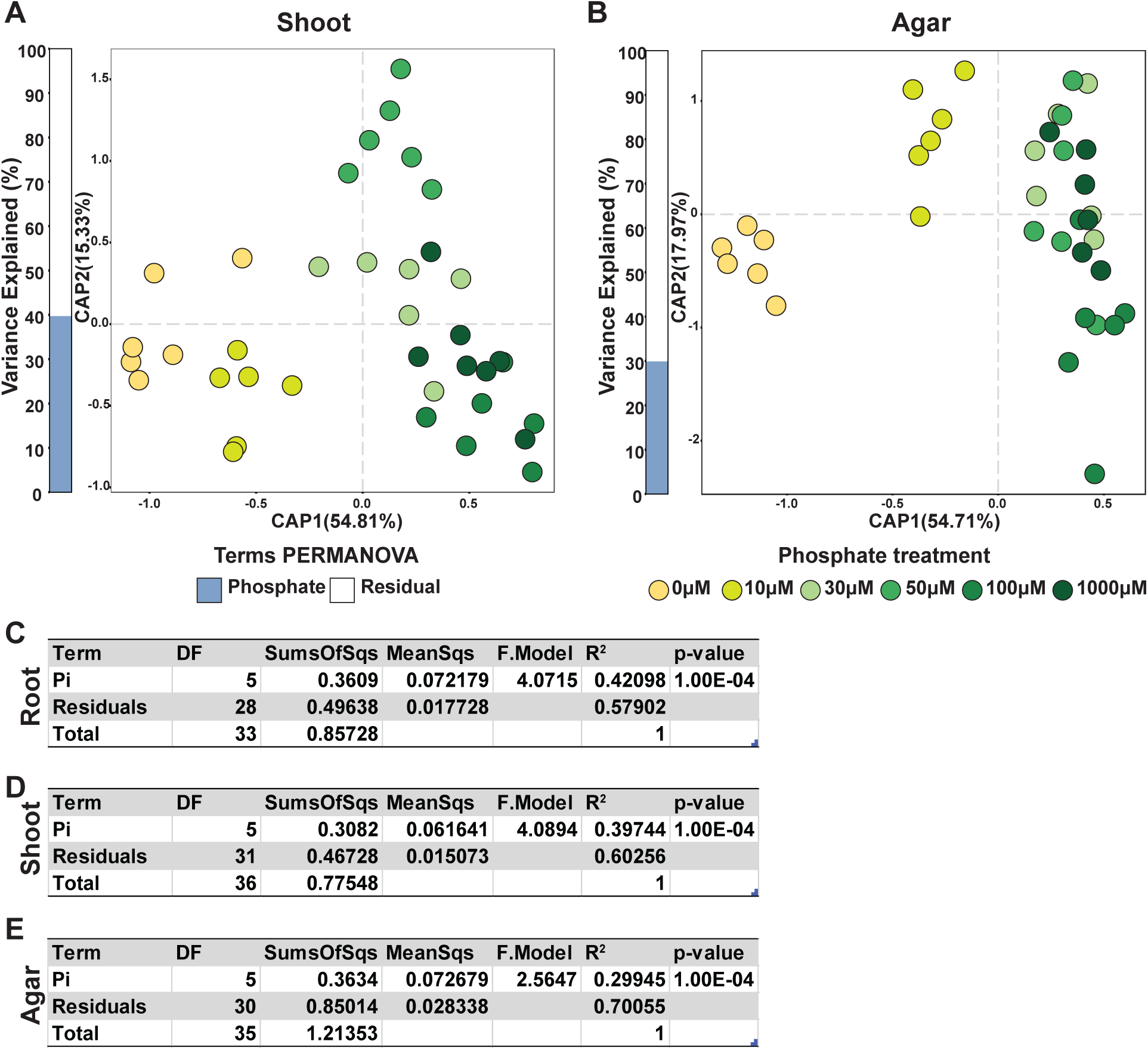
Bacterial synthetic community responds to the phosphate concentration in the media. **(A,B)** Canonical analysis of principal coordinates showing the influence of Pi concentration in the media on the bacterial communities in the **(A)** plant shoot and **(B)** agar. The bar graphs to the left of each plot depict the percentage of variability explained by statistically significant (p-value < 0.05) variables based on a PERMANOVA model (Materials and methods 4d). **(C,D,E)** PERMANOVA model results showing the influence of Pi concentration on the assembly of the bacterial community in **(C)** roots, **(D)** shoot and **(E)** agar.

**S7 Figure.**
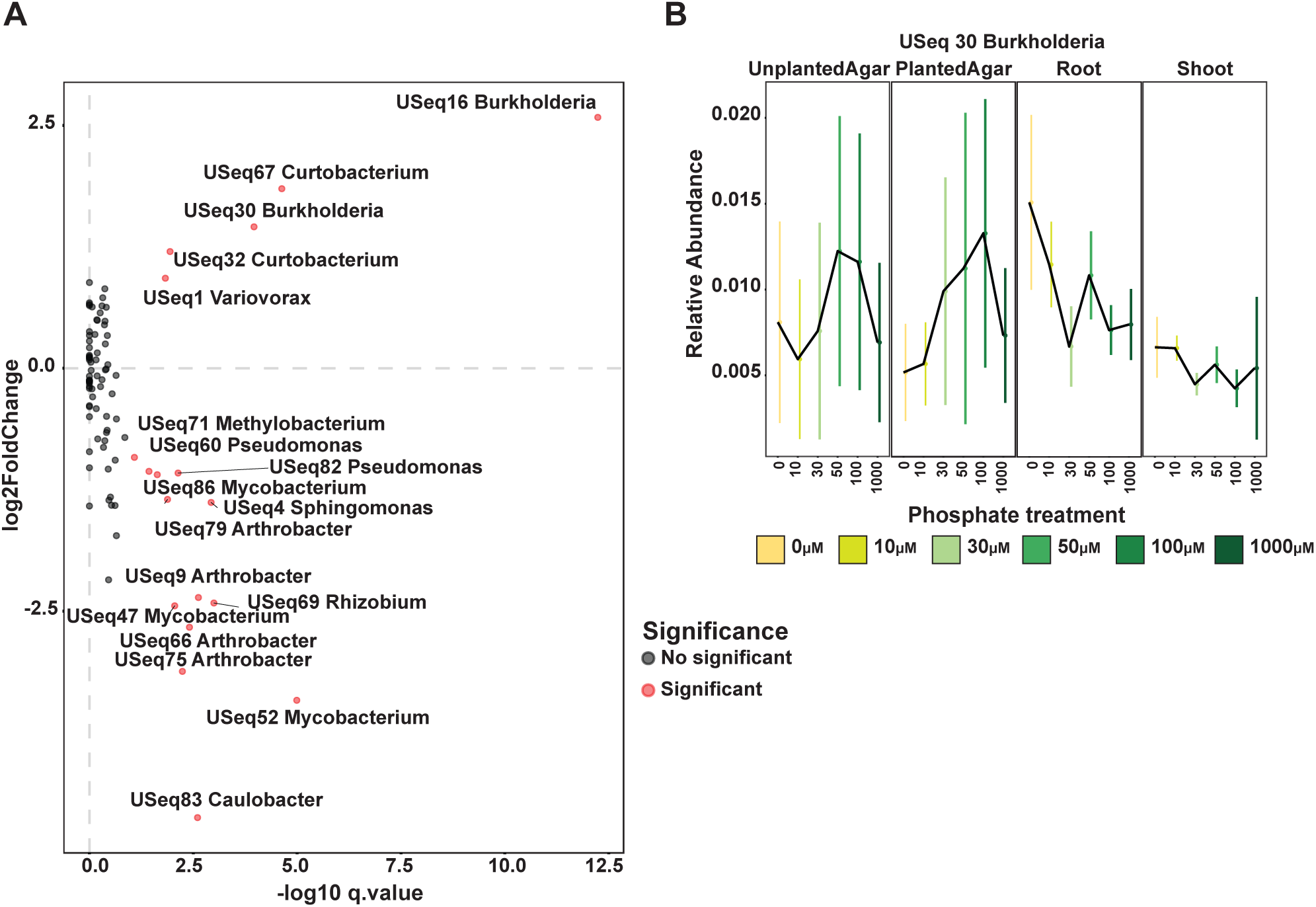
USeqs in the bacterial synthetic community displayed a strong Pi:fraction (shoot, root, agar) interaction. **(A)** Scatter plot (volcano plot) showing the results of the GLM interaction model between fraction and Pi concentration. The axes of the plot represent the output of the statistical test. The x-axis is the transformed q-value and the y-axis the log2 fold change. Each dot in the scatterplot represents a USeq. USeqs that showed a statistically significant fraction:Pi interaction are colored in red. USeqs genus and ID are displayed. The top right quadrant represents USeqs that are enriched in the plant tissues under low Pi conditions. **(B)** Relative abundance of Burkholderia Useq 30 that exhibits a statistically significant (q-value < 0.1) Pi-enrichment between the plant fractions and the agar fraction (Materials and methods 4d). The middle dot of each strip bar corresponds to the mean of that particular condition, the range of the strip bar corresponds to the 95% confidence interval of the mean.

## References

1. Desbrosses GJ, Stougaard J. Root Nodulation: A Paradigm for How Plant-Microbe Symbiosis Influences Host Developmental Pathways. Cell Host Microbe. 2011;10: 348–358. doi:10.1016/j.chom.2011.09.005

2. Hacquard S, Kracher B, Hiruma K, Münch PC, Garrido-Oter R, Thon MR, et al. Survival trade-offs in plant roots during colonization by closely related beneficial and pathogenic fungi. Nat Commun. 2016;7: 11362. doi:10.1038/ncomms11362

3. Dangl JL, Jones JDG. Plant pathogens and integrated defence responses to infection. Nature. 2001;411: 826–833. doi:10.1038/35081161

4. Castrillo G, Teixeira PJPL, Paredes SH, Law TF, de Lorenzo L, Feltcher ME, et al. Root microbiota drive direct integration of phosphate stress and immunity. Nature. 2017;543: 513–518. doi:10.1038/nature21417

5. Hacquard S, Spaepen S, Garrido-Oter R, Schulze-Lefert P. Interplay Between Innate Immunity and the Plant Microbiota. Annu Rev Phytopathol. 2017;55: 565–589. doi:10.1146/annurev-phyto-080516-035623

6. Haney CH, Samuel BS, Bush J, Ausubel FM, Gallo RL, Hooper L V., et al. Associations with rhizosphere bacteria can confer an adaptive advantage to plants. Nat Plants. 2015;1: 15051. doi:10.1038/nplants.2015.51

7. Pickles BJ, Wilhelm R, Asay AK, Hahn AS, Simard SW, Mohn WW. Transfer of ^13^ C between paired Douglas-fir seedlings reveals plant kinship effects and uptake of exudates by ectomycorrhizas. New Phytol. 2016;214: 400–411. doi:10.1111/nph.14325

8. Hernández M, Dumont MG, Yuan Q, Conrad R. Different bacterial populations associated with the roots and rhizosphere of rice incorporate plant-derived carbon. Appl Environ Microbiol. 2015;81: 2244–53. doi:10.1128/AEM.03209-14

9. Eissfeller V, Beyer F, Valtanen K, Hertel D, Maraun M, Polle A, et al. Incorporation of plant carbon and microbial nitrogen into the rhizosphere food web of beech and ash. Soil Biol Biochem. 2013;62: 76–81. doi:10.1016/j.soilbio.2013.03.002

10. Hu J, Wei Z, Friman V-P, Gu S-H, Wang X-F, Eisenhauer N, et al. Probiotic Diversity Enhances Rhizosphere Microbiome Function and Plant Disease Suppression. MBio. 2016;7: e01790–16. doi:10.1128/mBio.01790-16

11. Santhanam R, Luu VT, Weinhold A, Goldberg J, Oh Y, Baldwin IT. Native root-associated bacteria rescue a plant from a sudden-wilt disease that emerged during continuous cropping. Proc Natl Acad Sci U S A. 2015;112: E5013–20. doi:10.1073/pnas.1505765112

12. Mendes R, Kruijt M, de Bruijn I, Dekkers E, van der Voort M, Schneider JHM, et al. Deciphering the Rhizosphere Microbiome for Disease-Suppressive Bacteria. Science. 2011;332: 1097–1100. doi:10.1126/science.1203980

13. Shen Z, Ruan Y, Xue C, Zhong S, Li R, Shen Q. Soils naturally suppressive to banana Fusarium wilt disease harbor unique bacterial communities. Plant Soil. 2015;393: 21–33. doi:10.1007/s11104-015-2474-9

14. Cha J-Y, Han S, Hong H-J, Cho H, Kim D, Kwon Y, et al. Microbial and biochemical basis of a Fusarium wilt-suppressive soil. ISME J. 2016;10: 119–129. doi:10.1038/ismej.2015.95

15. Yuan Z, Druzhinina IS, Labbé J, Redman R, Qin Y, Rodriguez R, et al. Specialized Microbiome of a Halophyte and its Role in Helping Non-Host Plants to Withstand Salinity. Sci Rep. 2016;6: 32467. doi:10.1038/srep32467

16. Vargas L, Santa Brígida AB, Mota Filho JP, de Carvalho TG, Rojas CA, Vaneechoutte D, et al. Drought Tolerance Conferred to Sugarcane by Association with Gluconacetobacter diazotrophicus: A Transcriptomic View of Hormone Pathways. PLoS One. 2014;9: e114744. doi:10.1371/journal.pone.0114744

17. Hiruma K, Gerlach N, Sacristán S, Nakano RT, Hacquard S, Kracher B, et al. Root Endophyte Colletotrichum tofieldiae Confers Plant Fitness Benefits that Are Phosphate Status Dependent. Cell. 2016;165: 464–474. doi:10.1016/j.cell.2016.02.028

18. Vessey JK. Plant growth promoting rhizobacteria as biofertilizers. Plant Soil. 2003;255: 571–586. doi:10.1023/A:1026037216893

19. Fierer N, Jackson RB. The diversity and biogeography of soil bacterial communities. Proc Natl Acad Sci U S A. 2006;103: 626–31. doi:10.1073/pnas.0507535103

20. Lundberg DS, Lebeis SL, Paredes SH, Yourstone S, Gehring J, Malfatti S, et al. Defining the core Arabidopsis thaliana root microbiome. Nature. Nature Research; 2012;488: 86–90. doi:10.1038/nature11237

21. Yeoh YK, Paungfoo-Lonhienne C, Dennis PG, Robinson N, Ragan MA, Schmidt S, et al. The core root microbiome of sugarcanes cultivated under varying nitrogen fertilizer application. Environ Microbiol. 2016;18: 1338–1351. doi:10.1111/1462-2920.12925

22. Edwards J, Johnson C, Santos-medellín C, Lurie E, Kumar N. Structure, variation, and assembly of the root-associated microbiomes of rice. Proc Natl Acad Sci U S A. 2015;112: E911–E920. doi:10.1073/pnas.1414592112

23. Zhalnina K, Dias R, de Quadros PD, Davis-Richardson A, Camargo FAO, Clark IM, et al. Soil pH Determines Microbial Diversity and Composition in the Park Grass Experiment. Microb Ecol. 2015;69: 395–406. doi:10.1007/s00248-014-0530-2

24. Lauber CL, Hamady M, Knight R, Fierer N. Pyrosequencing-based assessment of soil pH as a predictor of soil bacterial community structure at the continental scale. Appl Environ Microbiol. 2009;75: 5111–20. doi:10.1128/AEM.00335-09

25. Griffiths RI, Thomson BC, James P, Bell T, Bailey M, Whiteley AS. The bacterial biogeography of British soils. Environ Microbiol. 2011;13: 1642–1654. doi:10.1111/j.1462-2920.2011.02480.x

26. Santos-Medellín C, Edwards J, Liechty Z, Nguyen B, Sundaresan V. Drought Stress Results in a Compartment-Specific Restructuring of the Rice Root-Associated Microbiomes. MBio. 2017;8: e00764–17. doi:10.1128/mBio.00764-17

27. Naylor D, DeGraaf S, Purdom E, Coleman-Derr D. Drought and host selection influence bacterial community dynamics in the grass root microbiome. ISME J. 2017; doi:10.1038/ismej.2017.118

28. Angel R, Soares MIM, Ungar ED, Gillor O. Biogeography of soil archaea and bacteria along a steep precipitation gradient. ISME J. 2010;4: 553–563. doi:10.1038/ismej.2009.136

29. Xu L, Naylor D, Dong Z, Simmons T, Pierroz G, Hixson KK, et al. Drought delays development of the sorghum root microbiome and enriches for monoderm bacteria. Proc Natl Acad Sci U S A. 2018;115: E4284–E4293. doi:10.1073/pnas.1717308115

30. Fitzpatrick CR, Copeland J, Wang PW, Guttman DS, Kotanen PM, Johnson MTJ. Assembly and ecological function of the root microbiome across angiosperm plant species. Proc Natl Acad Sci U S A. 2018;115: E1157–E1165. doi:10.1073/pnas.1717617115

31. Ramirez KS, Craine JM, Fierer N. Consistent effects of nitrogen amendments on soil microbial communities and processes across biomes. Glob Chang Biol. 2012;18: 1918–1927. doi:10.1111/j.1365-2486.2012.02639.x

32. Fierer N, Lauber CL, Ramirez KS, Zaneveld J, Bradford MA, Knight R. Comparative metagenomic, phylogenetic and physiological analyses of soil microbial communities across nitrogen gradients. ISME J. 2012;6: 1007–17. doi:10.1038/ismej.2011.159

33. Pantigoso HA, Manter DK, Vivanco JM. Phosphorus addition shifts the microbial community in the rhizosphere of blueberry (Vaccinium corymbosum L.). Rhizosphere. 2018;7: 1–7. doi:10.1016/J.RHISPH.2018.06.008

34. Yu P, Wang C, Baldauf JA, Tai H, Gutjahr C, Hochholdinger F, et al. Root type and soil phosphate determine the taxonomic landscape of colonizing fungi and the transcriptome of field-grown maize roots. New Phytol. Wiley/Blackwell (10.1111); 2018;217: 1240–1253. doi:10.1111/nph.14893

35. Leff JW, Jones SE, Prober SM, Barberán A, Borer ET, Firn JL, et al. Consistent responses of soil microbial communities to elevated nutrient inputs in grasslands across the globe. Proc Natl Acad Sci U S A. 2015;112: 10967–72. doi:10.1073/pnas.1508382112

36. Fabiańska I, Gerlach N, Almario J, Bucher M. Plant-mediated effects of soil phosphorus on the root-associated fungal microbiota in *Arabidopsis thaliana*. New Phytol. 2018;221: 2123–2137. doi:10.1111/nph.15538

37. Raghothama KG. PHOSPHATE ACQUISITION. Annu Rev Plant Physiol Plant Mol Biol. 1999;50: 665–693. doi:10.1146/annurev.arplant.50.1.665

38. Bustos R, Castrillo G, Linhares F, Puga MI, Rubio V, Pérez-Pérez J, et al. A Central Regulatory System Largely Controls Transcriptional Activation and Repression Responses to Phosphate Starvation in Arabidopsis. Ecker JR, editor. PLoS Genet. 2010;6: e1001102. doi:10.1371/journal.pgen.1001102

39. Gonzalez E, Solano R, Rubio V, Leyva A, Paz-Ares J. PHOSPHATE TRANSPORTER TRAFFIC FACILITATOR1 Is a Plant-Specific SEC12-Related Protein That Enables the Endoplasmic Reticulum Exit of a High-Affinity Phosphate Transporter in Arabidopsis. PLANT CELL ONLINE. 2005;17: 3500–3512. doi:10.1105/tpc.105.036640

40. Gransee A, Merbach W. Phosphorus dynamics in a long-term P fertilization trial on Luvic Phaeozem at Halle. J Plant Nutr Soil Sci. 2000;163: 353–357. doi:10.1002/1522-2624(200008)163:4<353::AID-JPLN353>3.0.CO;2-B

41. Robbins C, Thiergart T, Hacquard S, Garrido-Oter R, Gans W, Peiter E, et al. Root-Associated Bacterial and Fungal Community Profiles of Arabidopsis thaliana Are Robust Across Contrasting Soil P Levels. Phytobiomes. Phytobiomes; 2018;2: PBIOMES-09-17-0. doi:10.1094/PBIOMES-09-17-0042-R

42. Phillipson BA. PHOSPHATE TRANSPORTER TRAFFIC FACILITATOR1 Is a Plant-Specific SEC12-Related Protein That Enables the Endoplasmic Reticulum Exit of a High-Affinity Phosphate Transporter in Arabidopsis. Plant Cell Online. 2005;17: 3500–3512. doi:10.1105/tpc.105.036640

43. Knief C, Delmotte N, Chaffron S, Stark M, Innerebner G, Wassmann R, et al. Metaproteogenomic analysis of microbial communities in the phyllosphere and rhizosphere of rice. ISME J. 2012;6: 1378–1390. doi:10.1038/ismej.2011.192

44. Atamna-Ismaeel N, Finkel OM, Glaser F, Sharon I, Schneider R, Post AF, et al. Microbial rhodopsins on leaf surfaces of terrestrial plants. Environ Microbiol. 2012;14. doi:10.1111/j.1462-2920.2011.02554.x

45. Atamna-Ismaeel N, Finkel O, Glaser F, von Mering C, Vorholt JA, Koblížek M, et al. Bacterial anoxygenic photosynthesis on plant leaf surfaces. Environ Microbiol Rep. 2012;4: 209–216. doi:10.1111/j.1758-2229.2011.00323.x

46. Finkel OM, Delmont TO, Post AF, Belkin S. Metagenomic signatures of bacterial adaptation to life in the phyllosphere of a salt-secreting desert tree. Appl Environ Microbiol. 2016;82. doi:10.1128/AEM.00483-16

47. Levy A, Salas Gonzalez I, Mittelviefhaus M, Clingenpeel S, Herrera Paredes S, Miao J, et al. Genomic features of bacterial adaptation to plants. Nat Genet. 2018;50: 138–150. doi:10.1038/s41588-017-0012-9

48. Sieber M, Pita L, Weiland-Bräuer N, Dirksen P, Wang J, Mortzfeld B, et al. The Neutral Metaorganism. bioRxiv. 2018; 367243. doi:10.1101/367243

49. Kassen R, Buckling A, Bell G, Rainey PB. Diversity peaks at intermediate productivity in a laboratory microcosm. Nature. 2000;406: 508–512. doi:10.1038/35020060

50. Tilman D. Resource competition and community structure. Princeton University Press; 1982.

51. Herrera Paredes S, Gao T, Law TF, Finkel OM, Mucyn T, Teixeira PJPL, et al. Design of synthetic bacterial communities for predictable plant phenotypes. Kemen E, editor. PLOS Biol. 2018;16: e2003962. doi:10.1371/journal.pbio.2003962

52. Durán P, Thiergart T, Garrido-Oter R, Agler M, Kemen E, Schulze-Lefert P, et al. Microbial Interkingdom Interactions in Roots Promote Arabidopsis Survival. Cell. 2018;175: 973–983.e14. doi:10.1016/j.cell.2018.10.020

53. Baxter IR, Vitek O, Lahner B, Muthukumar B, Borghi M, Morrissey J, et al. The leaf ionome as a multivariable system to detect a plant’s physiological status. Proc Natl Acad Sci U S A. 2008;105: 12081–12086. doi:10.1073/pnas.0804175105

54. Merbach W, Garz J, Schliephake W, Stumpe H, Schmidt L. The long-term fertilization experiments in Halle (Saale), Germany — Introduction and survey. J Plant Nutr Soil Sci. 2000;163: 629–638. doi:10.1002/1522-2624(200012)163:6<629::AID-JPLN629>3.0.CO;2-P

55. Ames BN. [10] Assay of inorganic phosphate, total phosphate and phosphatases. Methods Enzymol. 1966;8: 115–118. doi:10.1016/0076-6879(66)08014-5

56. Logemann J, Schell J, Willmitzer L. Improved method for the isolation of RNA from plant tissues. Anal Biochem. 1987;163: 16–20. doi:10.1016/0003-2697(87)90086-8

57. Schindelin J, Arganda-Carreras I, Frise E, Kaynig V, Longair M, Pietzsch T, et al. Fiji: an open-source platform for biological-image analysis. Nat Methods. 2012;9: 676–682. doi:10.1038/nmeth.2019

58. Lundberg DS, Yourstone S, Mieczkowski P, Jones CD, Dangl JL. Practical innovations for high-throughput amplicon sequencing. Nat Methods. 2013;10: 999–1002. doi:10.1038/nmeth.2634

59. Gardes M, Bruns TD. ITS primers with enhanced specificity for basidiomycetes - application to the identification of mycorrhizae and rusts. Mol Ecol. 1993;2: 113–118. doi:10.1111/j.1365-294X.1993.tb00005.x

60. White TJ, Bruns, TD, Lee SB, Taylor JW. Amplification and direct sequencing of fungal ribosomal RNA genes for phylogenetics. PCR protocols: A Guide to Methods and Applications. 1990. pp. 315–322. doi:10.1016/B978-0-12-372180-8.50042-1

61. Taiyun Wei M, Taiyun Wei cre A, Simko aut V, Levy ctb M, Xie ctb Y, Jin ctb Y, et al. Package “corrplot”. 2017. Available from: https://cran.r-project.org/web/packages/corrplot/corrplot.pdf

62. Yourstone SM, Lundberg DS, Dangl JL, Jones CD. MT-Toolbox: improved amplicon sequencing using molecule tags. BMC Bioinformatics. 2014;15: 284. doi:10.1186/1471-2105-15-284

63. Joshi N, Fass J. Sickle: A sliding-window, adaptive, quality-based trimming tool for FastQ files (Version 1.33) [Software]. 2011. Available from: https://github.com/najoshi/sickle.

64. Callahan BJ, McMurdie PJ, Rosen MJ, Han AW, Johnson AJA, Holmes SP. DADA2: High-resolution sample inference from Illumina amplicon data. Nat Methods. 2016;13: 581–583. doi:10.1038/nmeth.3869

65. Caporaso JG, Kuczynski J, Stombaugh J, Bittinger K, Bushman FD, Costello EK, et al. QIIME allows analysis of high-throughput community sequencing data. Nat Methods. 2010;7: 335–336. doi:10.1038/nmeth.f.303

66. Edgar RC. Search and clustering orders of magnitude faster than BLAST. Bioinformatics. 2010;26: 2460–2461. doi:10.1093/bioinformatics/btq461

67. Deshpande V, Wang Q, Greenfield P, Charleston M, Porras-Alfaro A, Kuske CR, et al. Fungal identification using a Bayesian classifier and the Warcup training set of internal transcribed spacer sequences. Mycologia. 2016;108: 1–5. doi:10.3852/14-293

68. Salas Gonzalez I. isaisg/ohchibi: iskali. 2019; doi:10.5281/ZENODO.2593691

69. Oksanen J, Blanchet FG, Kindt R, Legendre P, Minchin PR, O’hara RB, et al. Package “vegan.” 2015. Available from: https://cran.r-project.org/web/packages/vegan/vegan.pdf

70. Lenth R, Singmann H, Love J, Buerkner P, Herve, M. Package “emmeans”. 2019; doi:10.1080/00031305.1980.10483031. Available from: https://cran.r-project.org/web/packages/emmeans/emmeans.pdf

71. Love MI, Huber W, Anders S. Moderated estimation of fold change and dispersion for RNA-seq data with DESeq2. Genome Biol. 2014;15: 550. doi:10.1186/s13059-014-0550-8

72. Letunic I, Bork P. Interactive tree of life (iTOL) v3: an online tool for the display and annotation of phylogenetic and other trees. Nucleic Acids Res. 2016;44: W242–W245. doi:10.1093/nar/gkw290

73. Mantel N. The detection of disease clustering and a generalized regression approach. Cancer Res. 1967;27: 209–20.

74. Edgar RC. UPARSE: highly accurate OTU sequences from microbial amplicon reads. Nat Methods. 2013;10: 996–998. doi:10.1038/nmeth.2604

75. Wang Q, Garrity GM, Tiedje JM, Cole JR. Naive Bayesian Classifier for Rapid Assignment of rRNA Sequences into the New Bacterial Taxonomy. Appl Environ Microbiol. 2007;73: 5261–5267. doi:10.1128/AEM.00062-07

76. DeSantis TZ, Hugenholtz P, Larsen N, Rojas M, Brodie EL, Keller K, et al. Greengenes, a chimera-checked 16S rRNA gene database and workbench compatible with ARB. Appl Environ Microbiol. 2006;72: 5069–72. doi:10.1128/AEM.03006-05

77. Wheeler TJ, Eddy SR. nhmmer: DNA homology search with profile HMMs. Bioinformatics. 2013;29: 2487–2489. doi:10.1093/bioinformatics/btt403

78. Katoh K, Standley DM. MAFFT multiple sequence alignment software version 7: improvements in performance and usability. Mol Biol Evol. 2013;30: 772–80. doi:10.1093/molbev/mst010

79. Capella-Gutiérrez S, Silla-Martínez JM, Gabaldón T. trimAl: a tool for automated alignment trimming in large-scale phylogenetic analyses. Bioinformatics. 2009;25: 1972–3. doi:10.1093/bioinformatics/btp348

80. Price MN, Dehal PS, Arkin AP. FastTree 2 – Approximately Maximum-Likelihood Trees for Large Alignments. PLoS One. 2010;5: e9490. doi:10.1371/journal.pone.0009490

81. Harvey, Pagel M. The comparative method in evolutionary biology. Oxford: Oxford University Press; 1991.

82. Revell LJ. phytools: an R package for phylogenetic comparative biology (and other things). Methods Ecol Evol. 2012;3: 217–223. doi:10.1111/j.2041-210X.2011.00169.x

83. Auguie B. Package “egg”. 2018. Available from: https://cran.r-project.org/web/packages/egg/egg.pdf

84. Andrews S. Babraham Bioinformatics - FastQC A Quality Control tool for High Throughput Sequence Data [Internet]. 2018. Available from: https://www.bioinformatics.babraham.ac.uk/projects/fastqc/. Cited 11 April 2019

85. Bolger AM, Lohse M, Usadel B. Trimmomatic: a flexible trimmer for Illumina sequence data. Bioinformatics. 2014;30: 2114–20. doi:10.1093/bioinformatics/btu170

86. Berardini TZ, Reiser L, Li D, Mezheritsky Y, Muller R, Strait E, et al. The arabidopsis information resource: Making and mining the “gold standard” annotated reference plant genome. genesis. 2015;53: 474–485. doi:10.1002/dvg.22877

87. Kim D, Langmead B, Salzberg SL. HISAT: a fast spliced aligner with low memory requirements. Nat Methods. 2015;12: 357–60. doi:10.1038/nmeth.3317

88. Liao Y, Smyth GK, Shi W. The Subread aligner: fast, accurate and scalable read mapping by seed-and-vote. Nucleic Acids Res. 2013;41: e108. doi:10.1093/nar/gkt214

89. Ewels P, Magnusson M, Lundin S, Käller M. MultiQC: summarize analysis results for multiple tools and samples in a single report. Bioinformatics. Narnia; 2016;32: 3047–3048. doi:10.1093/bioinformatics/btw354

90. Team RDC. R: A language and environment for statistical computing. 2011. Available from: https://www.r-project.org/

91. Yu G, Wang L-G, Han Y, He Q-Y. clusterProfiler: an R Package for Comparing Biological Themes Among Gene Clusters. O MICS. 2012;16: 284. doi:10.1089/OMI.2011.0118

